# Nanocatalytic activity of clean-surfaced, faceted nanocrystalline gold enhances remyelination in animal models of multiple sclerosis

**DOI:** 10.1101/712919

**Authors:** Andrew P. Robinson, Joanne Zhongyan Zhang, Haley E. Titus, Molly Karl, Mikhail Merzliakov, Adam R. Dorfman, Stephen Karlik, Michael G. Stewart, Richard K. Watt, Benjin D. Facer, Jon D. Facer, Noah D. Christian, Karen S. Ho, Michael T. Hotchkin, Mark G. Mortenson, Robert H. Miller, Stephen D. Miller

## Abstract

Development of pharmacotherapies that promote remyelination are a high priority for multiple sclerosis (MS) due to their potential for neuroprotection and restoration of function through repair of demyelinated lesions. A novel preparation of clean-surfaced, faceted gold nanocrystals demonstrated robust remyelinating activity in response to demyelinating agents in both chronic cuprizone and acute lysolecithin rodent animal models. Furthermore, oral delivery of gold nanocrystals improved motor functions of cuprizone-treated mice in both open field and kinematic gait studies. Gold nanocrystal treatment of oligodendrocyte precursor cells in culture resulted in oligodendrocyte maturation and expression of key myelin differentiation markers. Additional *in vitro* data demonstrated that these gold nanocrystals act via a novel energy metabolism pathway involving the enhancement of key indicators of aerobic glycolysis. In response to gold nanocrystals, co-cultured central nervous system cells exhibited elevated levels of the redox coenzyme nicotine adenine dinucleotide (NAD+), elevated total ATP levels, elevated extracellular lactate levels, and upregulation of myelin-synthesis related genes, collectively resulting in functional myelin generation. Based on these preclinical studies, clean-surfaced, faceted gold nanocrystals represent a novel remyelinating therapeutic for multiple sclerosis.

**One Sentence Summary:** Nanocatalytic activity of clean-surfaced, faceted gold nanocrystals results in robust remyelinating activity in demyelination animal models of multiple sclerosis.

## Introduction

Myelination is a complex process resulting in the wrapping of axons by oligodendrocyte (OL) membranes containing specialized proteins and lipids. The resulting myelin sheath facilitates axonal electrical conduction, which in turn supports a multitude of neuronal activities that coordinate motor, sensory, and higher order cognitive functions (*1*). Oligodendrocyte precursor cells (OPCs) differentiate into mature, myelinating OLs by following a differentiation program requiring input from both intrinsic and extrinsic signaling events. During active myelination, OLs synthesize on the order of 100,000 proteins per minute (*2*) and several thousand new lipid molecules per second (*3*), reflecting the significant energetic investment needed for biomass generation, and making this cell type among the most energetically demanding in the body. Dysregulated energy metabolism, particularly impacting OLs, has been postulated to play a central role in MS disease progression (*4, 5*). Limited remyelination occurs in multiple sclerosis (MS) lesions despite the presence of OPCs in or around lesion sites. These OPCs exhibit markers of metabolic stress, and, while capable of initiating a remyelination program, fail to do so (*6*). When cultured human OLs are metabolically stressed using low glucose media, OLs retract cellular processes and exhibit significantly reduced glycolytic activity, favoring cell survival at the expense of myelin stability (*7, 8*). By contrast, under favorable growth conditions, human OPCs and OLs preferentially utilize aerobic glycolysis over oxidative phosphorylation, resulting in the enhanced capacity to differentiate and myelinate axons (*7*). Aerobic glycolysis results in production of pyruvate and NADPH, the requisite precursors of myelin proteins and lipids. Preferential utilization of aerobic glycolysis has been observed in several developmental contexts in which large amounts of biomass, in the form of proteins and lipids, must be produced quickly (*9*). The enhancement of the glycolytic pathway has been proposed as a potential therapeutic target for remyelination in MS (*7*).

Unlike bulk gold which does not corrode and is generally recognized as chemically inert, gold at the nanoscale can be highly catalytic. Nanoparticulate gold catalysis is now widely used in many industrial applications, such as for the manufacture of propylene oxide (*10*), but only recently have biologically-relevant catalytic properties of gold nanoparticles been described (*11*). One of the key reactions catalyzed by gold nanoparticles is the oxidation of nicotinamide adenine dinucleotide hydride (NADH) to the critical energetic co-factor, NAD^+^ (*12*). NAD^+^ and NADH are not only metabolic sensors of cellular energy levels, but also serve as the essential redox couple for ATP-generating reactions, oxidative phosphorylation and glycolysis (*13*). In addition, NADH oxidation drives cellular respiratory and metabolic processes that play key roles in the energetically demanding process of myelination (*7, 8, 14*). Characterization of the catalysis of a redox reaction by a single gold nanoparticle using surface plasmon spectroscopy revealed that a gold 100 nm pentagonal bipyramid nanocrystal could accept approximately 4600 electrons per second, and reduce 65 O_2_ molecules per second, in the oxidation of ascorbic acid. Gold nanoparticles can thereby act as electron reservoirs with large capacity to donate or receive electrons when participating as redox reaction catalysts (*15*).

A novel electro-crystallization-based method was developed (US Patent 9,603,870) that results in suspensions of gold nanocrystals of 13 nm average diameter, termed CNM-Au8. This method resulted in clean, faceted surfaces on each nanocrystal, avoiding the potentially toxic deposition of organic residues on nanoparticle surfaces made using traditional synthesis methods (*16, 17*). CNM-Au8 exhibited significantly higher nanocatalytic activity than that of other commercially available comparator gold particles, and exhibited a clean toxicity profile in long-term six and nine-month rodent and canine chronic toxicology studies. Oral doses of CNM-Au8 were well tolerated in a First-in-Human Phase 1 clinical study. Here we describe the efficacy of CNM-Au8 to promote the robust remyelination of demyelinated axons and restore motor function in *in vivo* animal models of MS, induce differentiation of OPCs, and enhance activities of neurons and OLs through support of bioenergetic processes. Based on these results, CNM-Au8 is a viable therapeutic candidate for the treatment of remyelination failure in MS.

## Results

### Characterization of CNM-Au8

CNM-Au8 is a stable aqueous suspension of clean-surfaced nanocrystals consisting solely of gold (Au) atoms organized into crystals of highly faceted, geometrical shapes, which predominantly consist of hexagonal bi-pyramids, pentagonal bi-pyramids, octahedrons, and tetrahedrons as analyzed by transmission electron microscopy (TEM). Each Au nanocrystal has an approximate composition of up to 68,000 Au atoms per nanocrystal at the 13 nm median diameter, depending upon aspect ratio, with a corresponding molar mass of up to 1.3 x 10^4^ kDa per nanocrystal. The faceted structure of a single gold nanocrystal produced by electrocrystallization and imaged by transmission electron microscopy (TEM) is shown in Fig S1.

**Fig. S1:**
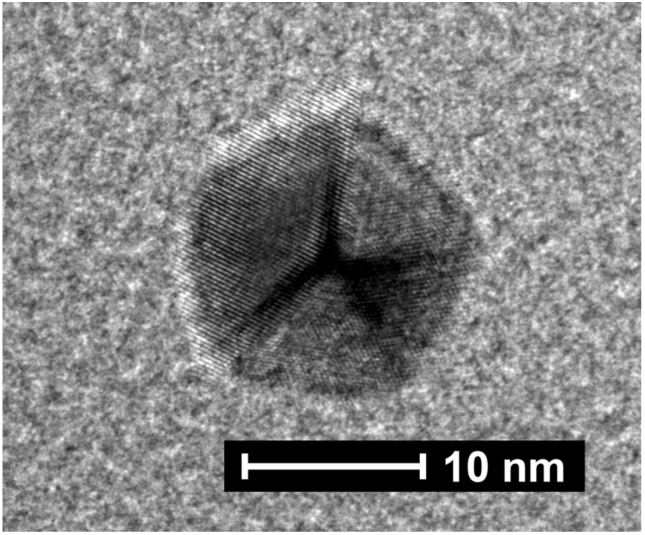
TEM image of a representative 13 nm pentagonal bipyramidal gold nanocrystal

Colloidal gold nanoparticles have previously been shown to catalyze the oxidation of NADH to NAD^+^ in a cell-free system (*12*). Utilizing the same UV-Vis spectrometry assay (*12*), we demonstrated that CNM-Au8 also efficiently catalyzed the oxidation of NADH to NAD^+^ (Fig. 1A-D). Fig.1A shows the time course of the change in absorbance spectra of NADH (seen by a diminishing peak at 339 nm) with a concomitant increase in the NAD^+^ peak at 259 nm. The control, conducted and sampled over the same time period without the addition of CNM-Au8, showed no spectral changes in NADH absorbance, demonstrating that NADH alone was stable and did not oxidize under these conditions without the presence of CNM-Au8 (Fig. 1C, black curve). Furthermore, the oxidation of NADH by CNM-Au8 was dose-dependent; Fig. 1B shows NADH absorbance at 339 nm as a function of time for four different reaction concentrations of CNM-Au8 (6.6, 12.4, 23.4, and 46.8 µg/mL).

**Figure 1:**
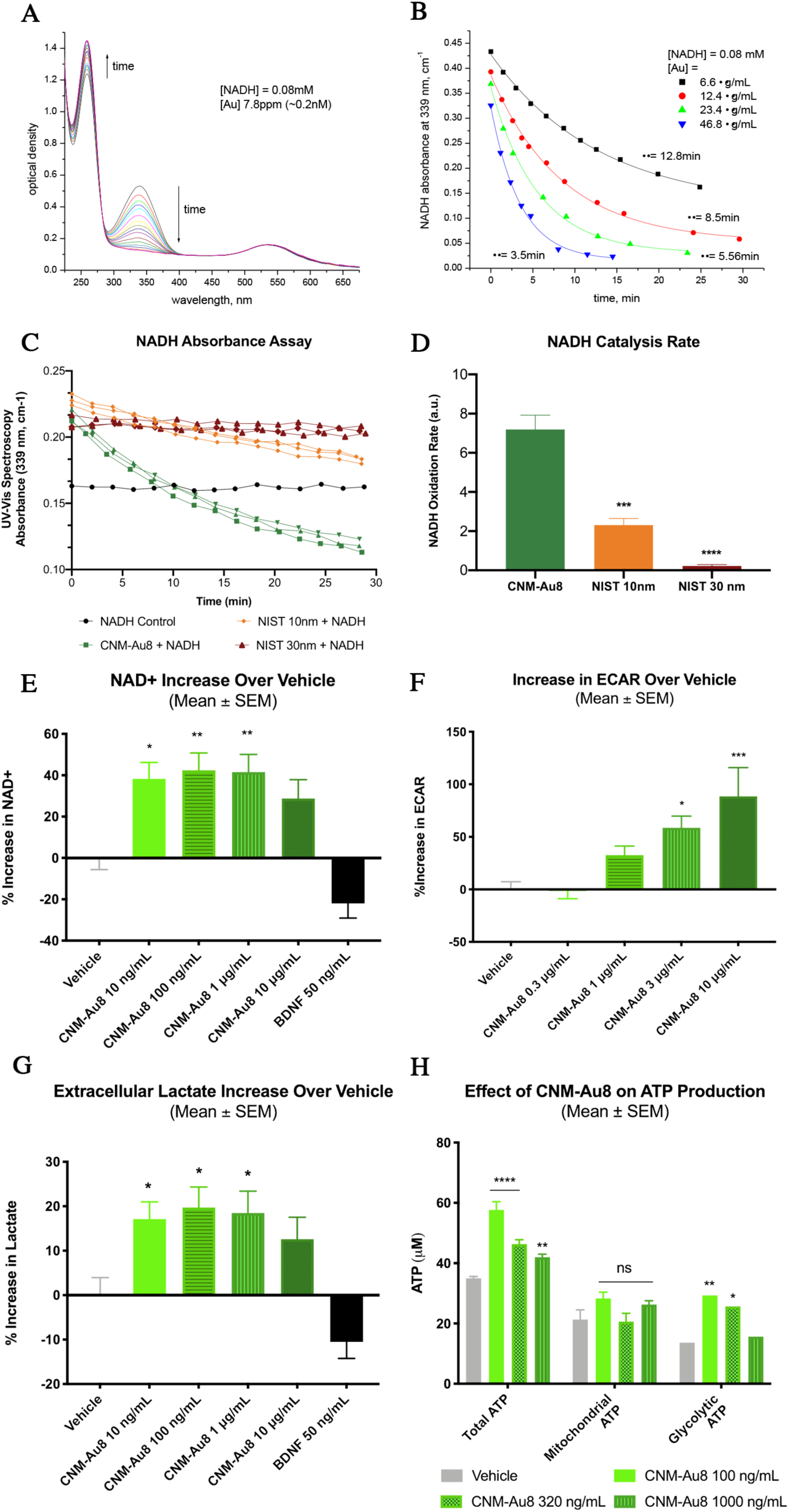
Nanocatalysis by CNM-Au8 enhances cellular bioenergetics. A, Changes in absorbance peaks of NADH (339 nm) and NAD+ (259 nm) demonstrate the conversion of NADH to NAD+ with time in the presence of 0.2 nM CNM-Au8. Shown is an overlay of UV-Vis spectra taken at approximately 30-s intervals; total elapsed time of 60 minutes. Starting concentration of NADH at t=0 was 0.08 mM. B, Dose-dependent catalytic activity of CNM-Au8 (6.6 g/mL black squares, 12.4 g/mL red circles, 23.4 g/mL green triangles, 46.8 g/mL blue triangles) on the oxidation of NADH as measured by change in absorbance of NADH over time. C, Catalytic activity of CNM-Au8 compared to two NIST standards, NIST 10 nm (orange) and NIST 30 nm (red), using the same starting concentrations: 3.4 ppm Au added to 26 μM NADH in 5.7 mM NaHCO_3_. No gold control (black) shows stability of 26 μM NADH during the same time frame. D, Initial rates of catalysis for CNM-Au8, NIST 10 nm, and NIST 30 nm, calculated from curves shown in B. E, Effect of CNM-Au8 treatment on NAD+ levels, expressed as percent change over vehicle, in primary mesencephalic cultures compared to vehicle (gray), or BDNF control. F, Extracellular acidification rate (ECAR) of purified murine OLs in response to CNM-Au8 in the first three minutes following glucose challenge as measured in the Seahorse flux analyzer, expressed as percent change over vehicle. G, Extracellular lactate levels in media of primary rat mesencephalic cultures after 48h treatment with CNM-Au8, expressed as percent change over vehicle. H, Total, mitochondrial, and glycolytic intracellular ATP levels from human OL M03.13 cells treated with vehicle (gray) or CNM-Au8 (green). D-G, One-way ANOVA, corrected for multiple comparisons. H, Two-way ANOVA. Quantities shown are group means +/-SEM. * p < 0.05; ** p < 0.01; *** p < 0.001; **** p < 0.0001.

To determine whether catalytic rates differed between electro-crystallization-derived CNM-Au8 and gold nanoparticles of similar diameter synthesized using the commonly used citrate reduction of chloroauric acid method (*18, 19*), we used two reference standards of gold nanoparticles from the U.S. National Institute of Standards and Technology (NIST) that were 10 nm and 30 nm in diameter, respectively at equivalent concentrations to CNM-Au8. Under the same reaction conditions, the nanocatalytic activity of CNM-Au8 significantly exceeded the catalytic activity of the citrate-reduced gold nanoparticle standards of both diameters by over 3-fold (Fig. 1C, D).

An effect on the total intracellular NAD+ pool due to treatment with CNM-Au8 was also observed in living cells. Primary co-cultures of neural and glial cells from the mesencephalon of day 15 rat embryos were exposed to CNM-Au8, brain derived neurotrophic growth factor (BDNF, a control for neuroprotection), or vehicle for 36 hours. Total cellular NAD^+^ concentrations in these treated culture lysates were quantified using a commercial luciferase-based assay kit. Figure 1E demonstrates that significantly elevated NAD+ intracellular levels are detected in cells following treatment with CNM-Au8.

The effects of CNM-Au8 to increase the net pool of NAD^+^ across mixed CNS cell types suggested that CNM-Au8 treatment could support the glycolytic and anabolic processes key to OL myelination. To measure elevated glycolytic activity, the Seahorse Flux Analyzer (Agilent) extracellular acidification rate (ECAR) assay was used to measure the effect of CNM-Au8 treatment on the glycolytic activity in OLs immediately following glucose challenge. Figure 1F shows a CNM-Au8 dose-dependent increase in glycolytic activity in OLs immediately following glucose challenge in the Seahorse assay. Lactate, produced as a result of elevated glycolytic activity, is the primary cause of acidification of media. Figure 1G shows that CNM-Au8 treatment of primary neuronal-glial co-cultures elevated the levels of extracellular lactate. Further, an increase in non-mitochondrial ATP production would be expected to result from enhanced glycolytic activity. To demonstrate CNM-Au8’s effect on non-mitochondrial ATP levels, an immortalized oligodendrocyte human cell line, MO3.13 cells, were seeded into two plates at the same density and cultured with CNM-Au8 or vehicle for 72 hours. One plate was additionally treated with a cocktail of mitochondrial inhibitors (oligomycin, rotenone, and antimycin A) for two hours prior to lysis to block all mitochondrial ATP production, and the other plate served as the non-treated control. Cells were then immediately lysed in lysis buffer containing luciferin and total ATP was measured from both plates using a luciferase assay. ATP derived from glycolytic activity significantly contributed to the enhanced levels of total ATP in CNM-Au8-treated human OLs compared to the vehicle controls (Fig. 1H).

### Remyelination by nanocrystalline gold using the cuprizone in vivo model of demyelination

The copper-chelating OL toxin cuprizone selectively induces the apoptosis of mature OLs in the CNS of mice following a standard five-week toxin treatment protocol (*20*). With five weeks of cuprizone feeding, maximal demyelination of axons in the corpus callosum can be achieved (*20*). This model is commonly used as a model for MS as well as other white matter degenerative diseases.

To determine whether CNM-Au8 had a therapeutic effect on mice exposed to cuprizone, CNM-Au8 was delivered to randomized groups of mice (N=15 per group) by gavage (10 mg/kg/day) (Groups 1-5) or was provided as their drinking water *ad libitum* (Groups 6 and 7). A prophylactic treatment arm with CNM-Au8 was investigated by providing CNM-Au8 to Groups 4 and 6 concomitant with the start of CPZ treatment, while a therapeutic treatment arm with CNM-Au8 was investigated by starting CNM-Au8 dosing two weeks after cuprizone administration had been initiated (Groups 5 and 7). Group 1 served as a non-cuprizone-treated vehicle control (sham), with vehicle provided by gavage daily. To assess the time-course of cuprizone damage, Group 2 was started on cuprizone and vehicle by gavage, then sacrificed after two weeks to verify the ongoing cuprizone insult. Group 3 was a cuprizone and vehicle control that was sacrificed at the end of five weeks when maximal myelin damage was expected to have taken place (*20*). A schematic for the study design is shown in Fig. 2A.

**Figure 2:**
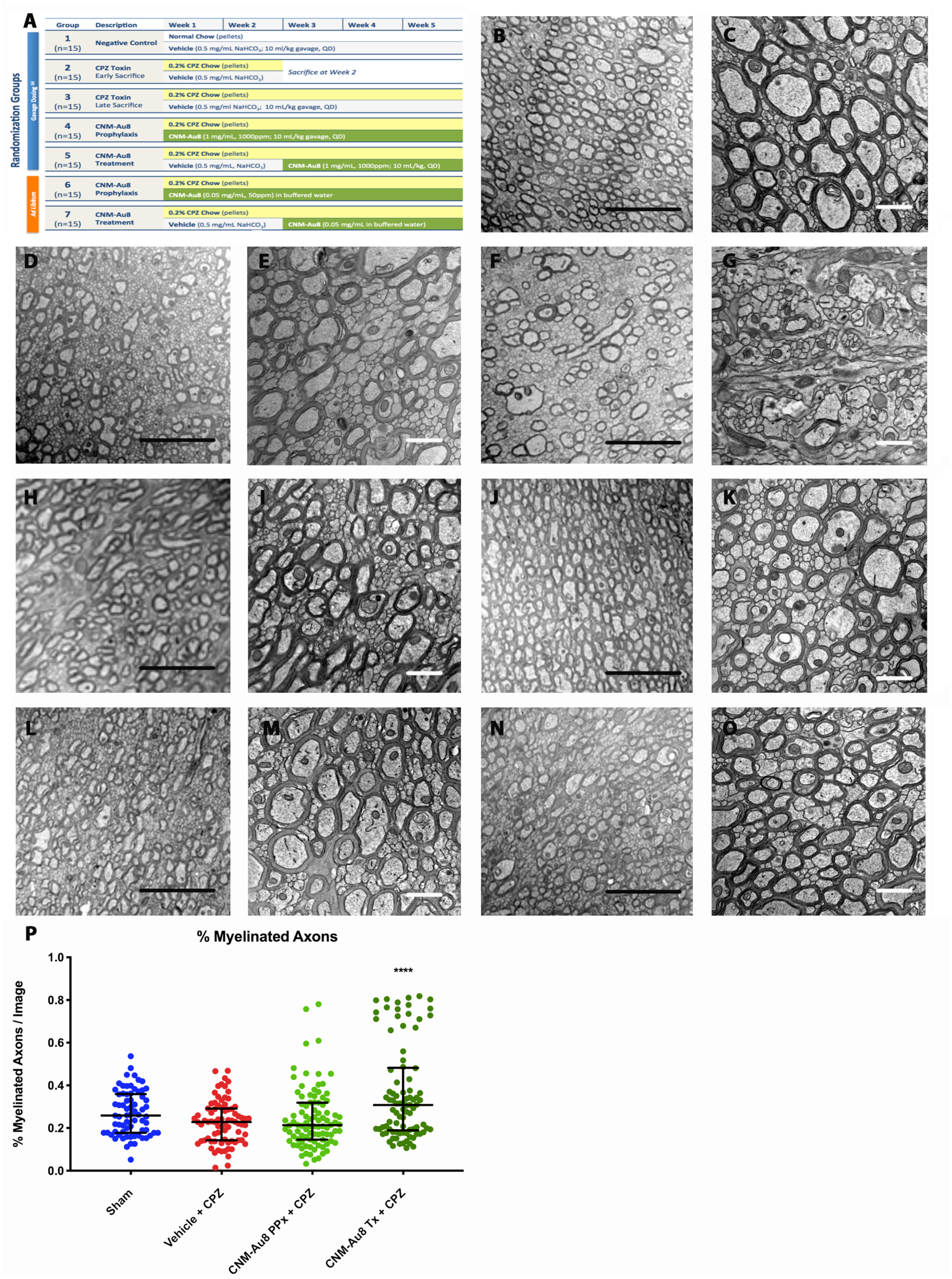
Effect of CNM-Au8 in an in vivo cuprizone model of demyelination. A. Experimental design schematic. Groups 1-5 were dosed by gavage and Groups 6 and 7 were dosed ad libitum. For Groups 4 and 6, administration of CNM-Au8 began at Week 1, concomitant with cuprizone administration (prophylactic arm); for Groups 5 and 7, administration of CNM-Au8 began at Week 3 (treatment arm). B-O, Representative TEM images of corpus callosum axons in cross section from sham-treated Group 1 animals (B,C), cuprizone-treated animals sacrificed at Week 2 (D,E), cuprizone-treated animals sacrificed at Week 5 (F,G), cuprizone plus gavage-dosed CNM-Au8 treated animals from the prophylactic arm (H,I), cuprizone plus gavage-dosed CNM-Au8 treated animals from the therapeutic arm (J,K), cuprizone plus ad libitum-dosed CNM-Au8 treated animals from the prophylactic arm (L,M),and cuprizone plus ad libitum-dosed CNM-Au8 treated animals from the therapeutic arm (N,O). Magnifications were 4000x (B,D,F,H,J,L,N) and 16,000x (C,E,G,I,K,M,O). Black bar = 5 μM; white bar = 1 μM. P, Quantitation of the percent myelinated axons in 16,000x TEM images by treatment group. Error bars show median and interquartile ranges of each group. **** p < 0.0001, one-way ANOVA corrected for multiple comparisons.

TEM images of the corpus callosum of animals from each group were analyzed both qualitatively by an expert independent pathologist and quantitatively in a blinded fashion. After two weeks of cuprizone treatment, significant changes in myelin morphology were noted. In comparison to the ordered array of well-packed and ovoid-shaped, myelinated axons in the vehicle-treated control (Fig. 2B, C), cuprizone feeding caused extensive CNS damage: the tissue appeared disorganized, frequent delamination of lamellae was apparent, axons were irregularly-shaped, and patches of demyelinated axons were observed within areas of normal-appearing myelination (Fig. 2D-G). By contrast, in cuprizone-fed animals treated with CNM-Au8 (Fig. 2H-O), there were extensive areas of normal myelin observed, as well as very few swollen myelin sheaths and a reduced incidence of delamination.

To quantitate the changes in remyelination that could be attributed to CNM-Au8 treatment, ImageJ was used to count the number of myelinated and unmyelinated axons in each image (N=587 images; average of 84 images per treatment group) to calculate the percentage of myelinated axons per image. As expected, cuprizone treatment reduced the percentage of myelinated axons compared to sham treated animals (Fig. 2P, p = 0.0221, two-tailed Mann-Whitney test). Striking recovery of myelinated axons was observed for therapeutically treated animals who were dosed with CNM-Au8 by gavage (Fig. 2P, dark green scatterplot group). An increase in myelinated axons per image was also observed for animals in the prophylactic arm with CNM-Au8 administered by gavage; however, this trend did not reach statistical significance when corrections for multiple comparisons were applied (Fig. 2P, light green scatterplot). Animals dosed with CNM-Au8 *ad libitum* also displayed improved myelination in TEM images (Fig. 2L-O), however, quantitation of myelinated axons per image did not reach statistical significance (not shown).

These results corroborated those of an earlier pilot cuprizone experiment in which four groups of mice, randomized by body weight (N=4 per group), were treated with vehicle, cuprizone, CNM-Au8 alone, or CNM-Au8 and cuprizone (Fig. S2A). Higher numbers of myelin-wrapped axons in CNM-Au8 treated animals were readily seen by TEM (Fig. S2B, Group 4 vs. Group 2). Using gratio analysis, calculated as the ratio between the inner myelin circumference divided by the outer myelin circumference, a distinct population of thinly myelinated axons can be observed in the cuprizone-treated animals’ g-ratios, which is not evident in animals treated with both CNM-Au8 and cuprizone (Fig. S2C, purple scatter plot vs. red scatter plot). Higher levels of myelin proteolipid protein (PLP) expression, a marker of mature OL myelin production, were observed in coronal slices of cuprizone-administered animals treated with CNM-Au8 as compared to vehicle controls (Fig. S2D). Taken together, these demyelinating animal model studies indicated that CNM-Au8 treatment consistently resulted in higher levels of myelinated axons compared to vehicle treated controls.

**Figure S2:**
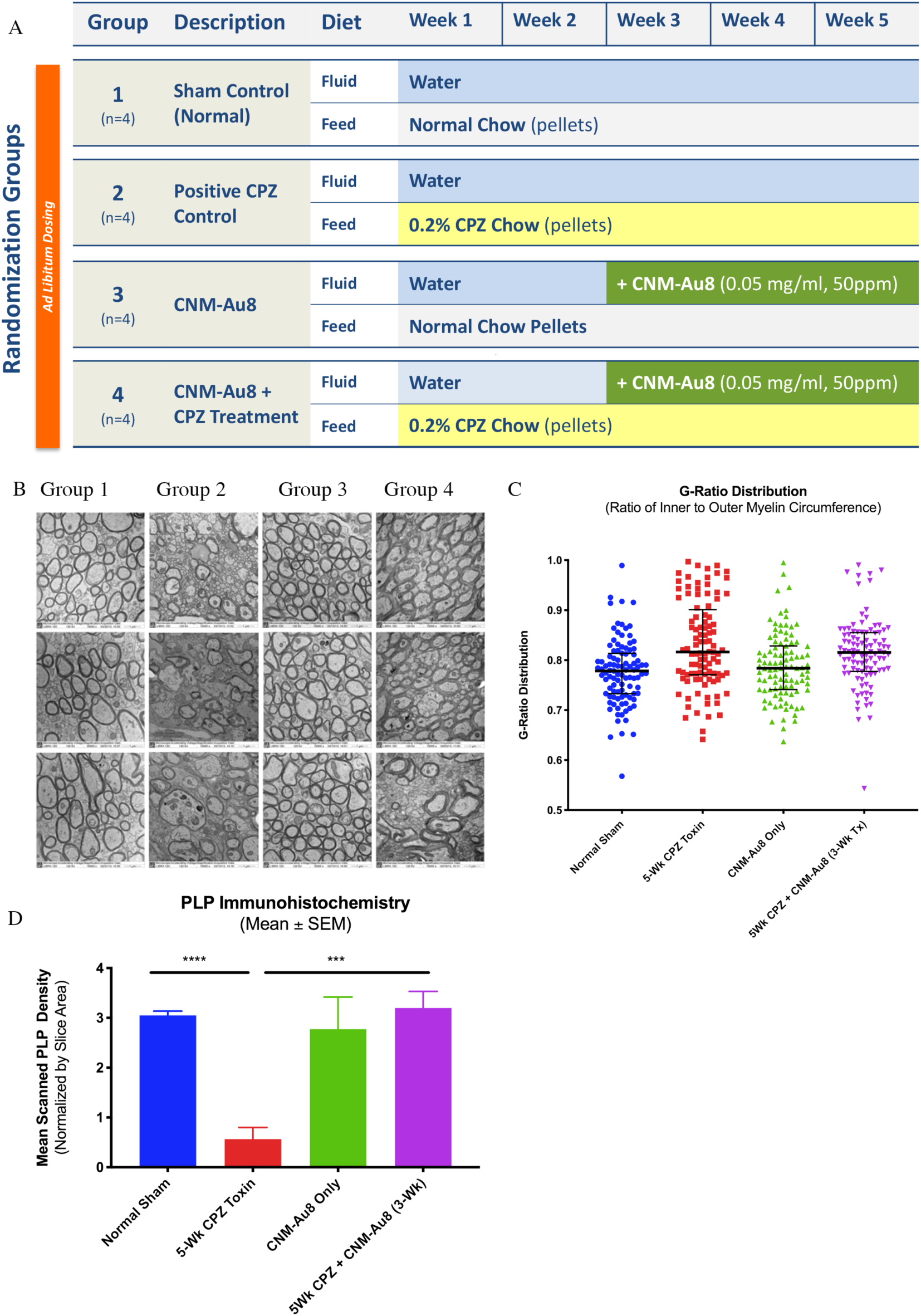
Evidence of remyelination by CNM-Au8 in a pilot in vivo cuprizone mouse model. A, Experimental design schematic. B, Three representative TEM images from the corpus callosum of mice from each group (16,000x magnification). C, G-ratio distribution scatter plots of 100 randomly selected axons from images of corpus callosum sections of one animal from each group, assessed in blinded fashion. Bars show the median with interquartile ranges. Smaller g-ratios indicate thicker myelin sheaths. D, Quantitation of anti-PLP staining of coronal sections of whole brain of one animal from each group, normalized by hematoxylin-stained area. *** p value < 0.001 using two-tailed Student’s t-test comparison to 5-wk cuprizone toxin (Group 2).

**Figure S3:**
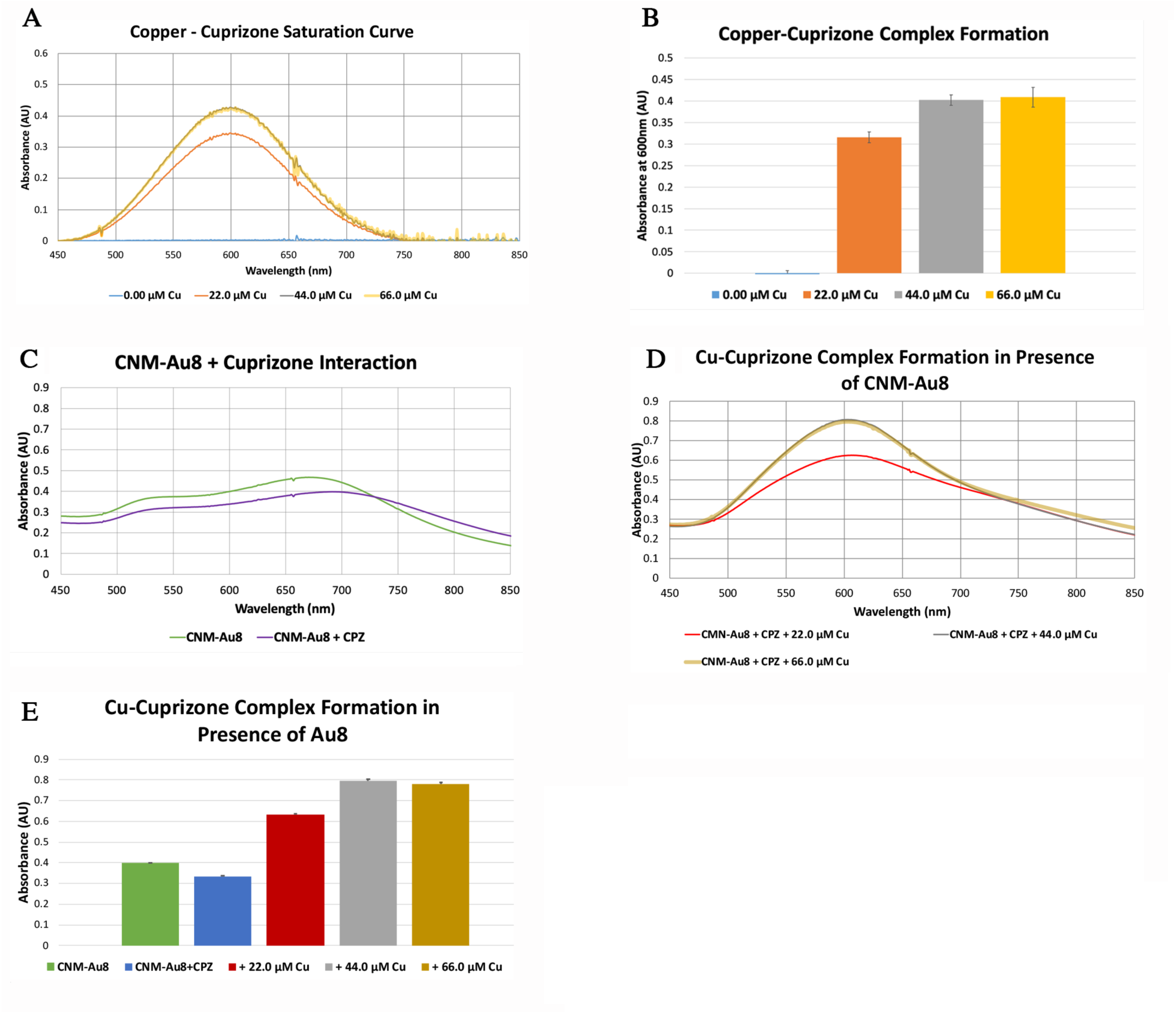
CNM-Au8 does not inhibit cuprizone from chelating copper. A, Copper-cuprizone complexes have a broad absorbance peak at 600 nm, observed upon the addition of varying amounts of Cu(II) sulfate to 88.0 μM cuprizone. Cuprizone becomes saturated on the addition of 44.0 μM Cu (gray), which is superimposed by the curve of 66.0 μM Cu (yellow). B, Quantitation of the average absorbance of the Cu-cuprizone complex at 600 nm (shown in A) for triplicate samples. C, Minimal changes in the UV-Vis absorbance profile of 26 μg/mL CNM-Au8 (∼2 μM) in the presence (purple) or absence (green) of 88.0 μM cuprizone indicates a lack of binding interaction between CNM-Au8 and cuprizone. The slight spectral shift may indicate possible minor surface interactions. D,E, CNM-Au8 does not interfere with Cu-cuprizone binding. Saturation of 88.0 μM cuprizone occurs with the addition of 44.0 uM copper (gray) in the presence of 26 μg/mL CNM-Au8. E, Quantitation of the average absorbance of the Cu-cuprizone complex in the presence of 26 μg/mL CNM-Au8 at 600 nm (shown in D) for triplicate samples. Saturation of cuprizone occurs with the addition of 44.0 uM copper (gray).

### Remyelination by CNM-Au8 does not occur via inactivation of cuprizone

Two possibilities could explain the actions of CNM-Au8 to improve myelin content in the brains of cuprizone treated animals. CNM-Au8 could act by protecting OLs from cuprizone-induced apoptosis, or CNM-Au8 could act by stimulating the differentiation of OPCs to mature and remyelinate. To distinguish between these two possibilities, we conducted the following studies. The first was to determine whether CNM-Au8 binds and sequesters cuprizone, or blocks its copper binding ability, in such a way that CNM-Au8 inactivated cuprizone. To detect alterations in cuprizone’s ability to bind copper in the presence of CNM-Au8, UV-Vis spectroscopy was used to monitor binding activities of copper to cuprizone in the presence and absence of CNM-Au8. Once it was established that the sequestration of copper by cuprizone was unaffected by the presence of CNM-Au8 (Fig. S3), we then carried out an additional *in vivo* cuprizone study, in which cuprizone was fed to mice for five weeks to achieve maximal demyelination, only after which CNM-Au8 or vehicle were provided. We then assessed the animals for remyelination during the period following the cessation of cuprizone dosing (Fig. 3).

**Figure 3:**
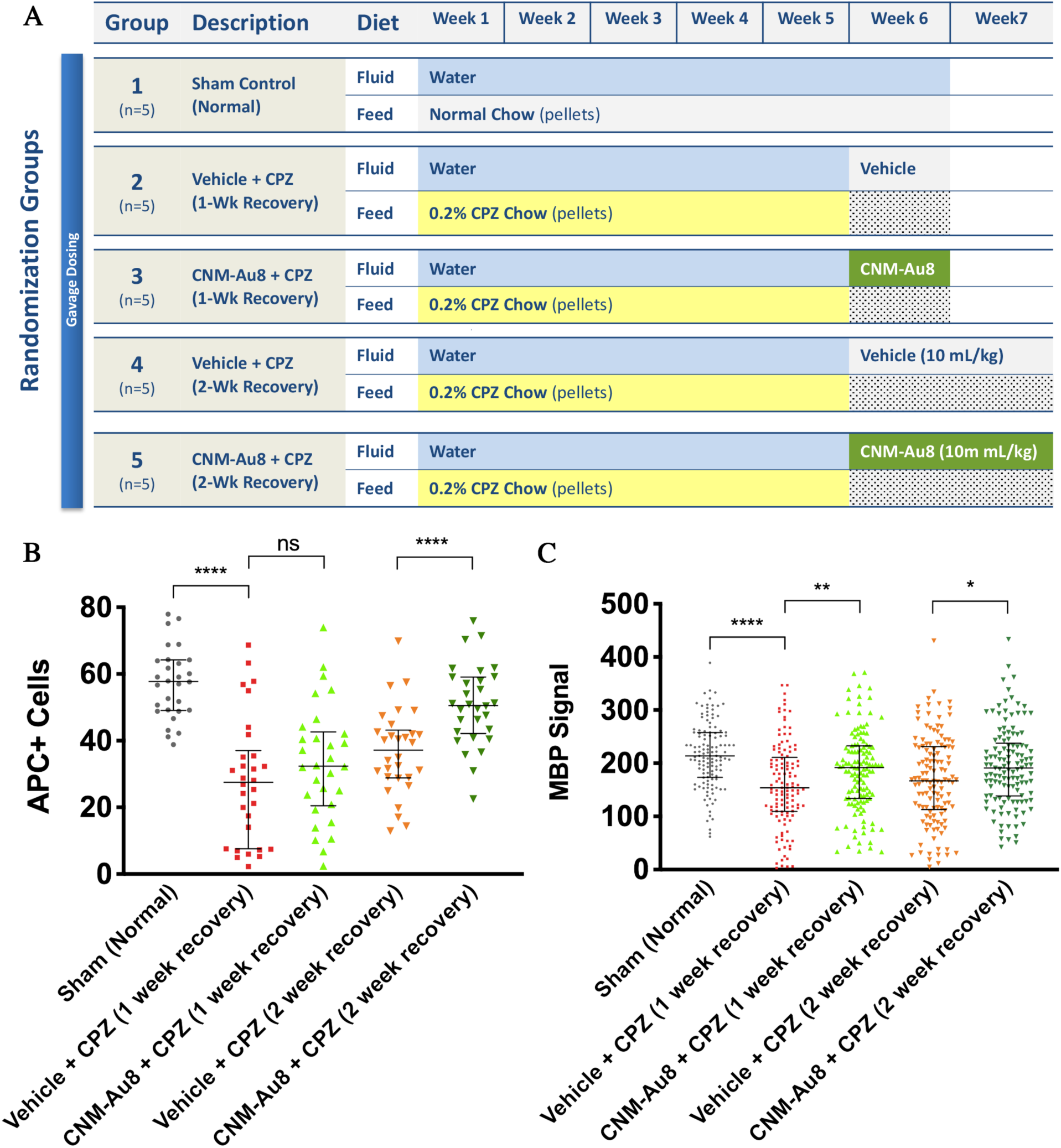
CNM-Au8 promotes oligodendrocyte maturation when administered post-cuprizone treatment. A, Experimental design schematic. B, Quantitation of the number of APC positive cells by immunohistochemical staining of coronal sections from each treatment group. C, Quantitation of the area of anti-MBP immunohistochemical staining of coronal sections from each treatment group. B,C, error bars show median and interquartile range.

For the first study, we established a saturation curve for copper binding to cuprizone. The interaction was characterized by an absorbance peak at 600 nm (Fig. S3A, B). A similar absorbance curve generated by a mixture of CNM-Au8 with cuprizone indicated an initial redshift in the absorbance curve immediately upon cuprizone addition (Fig. S3C), suggesting a weak surface interaction of CNM-Au8 with cuprizone. However, the absorbance peak of the saturation curve for copper and cuprizone in the presence of CNM-Au8 remained unaltered at 600nm, demonstrating the uninhibited interaction of cuprizone with copper in the presence of CNM-Au8 (Fig. S3D,E). Furthermore, when the background signal due to CNM-Au8 alone was subtracted, the absorption spectrum of copper-cuprizone complexes was the same as that of copper-cuprizone without CNM-Au8, thus demonstrating that the presence of CNM-Au8 did not affect the sequestration of copper by cuprizone (Fig. S3D,E).

Next, we designed an *in vivo* study in which cuprizone demyelination was carried out for five weeks (N=5 per group), then cuprizone was discontinued and treatment with CNM-Au8 or vehicle commenced and continued for the following week or two weeks (Fig. 3A). This experimental design allowed for the assessment of efficacy of CNM-Au8 after cuprizone was withdrawn. Figures 3B and C show that CNM-Au8 acts to stimulate myelin marker expression in OLs after cuprizone damage had taken place. The number of mature, myelin-producing OLs was assessed by antibody staining coronal sections from each mouse using anti-APC (CC1), a marker of mature OL cell bodies, and with anti-MBP, a marker of myelin-producing OLs, at the end of either one or two weeks of treatment with CNM-Au8 compared with vehicle controls. As shown in Fig.3B, the animals treated with CNM-Au8 for two weeks following cuprizone insult show an increase in the number of APC-positive cells in comparison with vehicle-treated control, which reached statistical significance for the two-week, CNM-Au8-treated group. This is supported by MBP immunohistochemistry results shown in Fig. 3C, in which an increase in MBP staining in CNM-Au8-treated mice over their respective vehicle controls is observed.

### Remyelination by CNM-Au8 does not occur by direct protection of mature OLs from cuprizone-mediated apoptosis

Cuprizone preferentially induces the apoptosis of mature OLs over OPCs and other neural cell types (*20*). To determine whether CNM-Au8 acts to protect mature OLs from apoptosis induced by cuprizone, a study was conducted in which mice were simultaneously treated with cuprizone and CNM-Au8 or vehicle control for two weeks during the initial onset of cuprizone OL toxicity, after which the animals were sacrificed and coronal brain sections were stained for mature OLs using anti-APC and anti-MBP antibodies (Fig. S4). Early assessment of the effects of cuprizone during this two-week period allowed us to detect a potential reduction in mature OL numbers before OPCs could differentiate and replenish the mature OL population.

After two weeks of cuprizone treatment, the number of OLs marked by anti-APC and MBP was reduced in all coronal sections, and this reduction was not improved by concurrent treatment of CNM-Au8 or prophylactic treatment of CNM-Au8 starting one week prior to cuprizone treatment (Fig. S4). This observed reduction in mature OL numbers and MBP expression, despite the administration of CNM-Au8 in cuprizone-fed animals, indicates that CNM-Au8 does not protect mature OLs from cuprizone induced toxicity. Recovery of myelin following CNM-Au8 treatment in cuprizone-treated animals is therefore not due to the improved survival of mature OLs, but more likely due to enhanced differentiation of OPCs.

**Figure S4.**
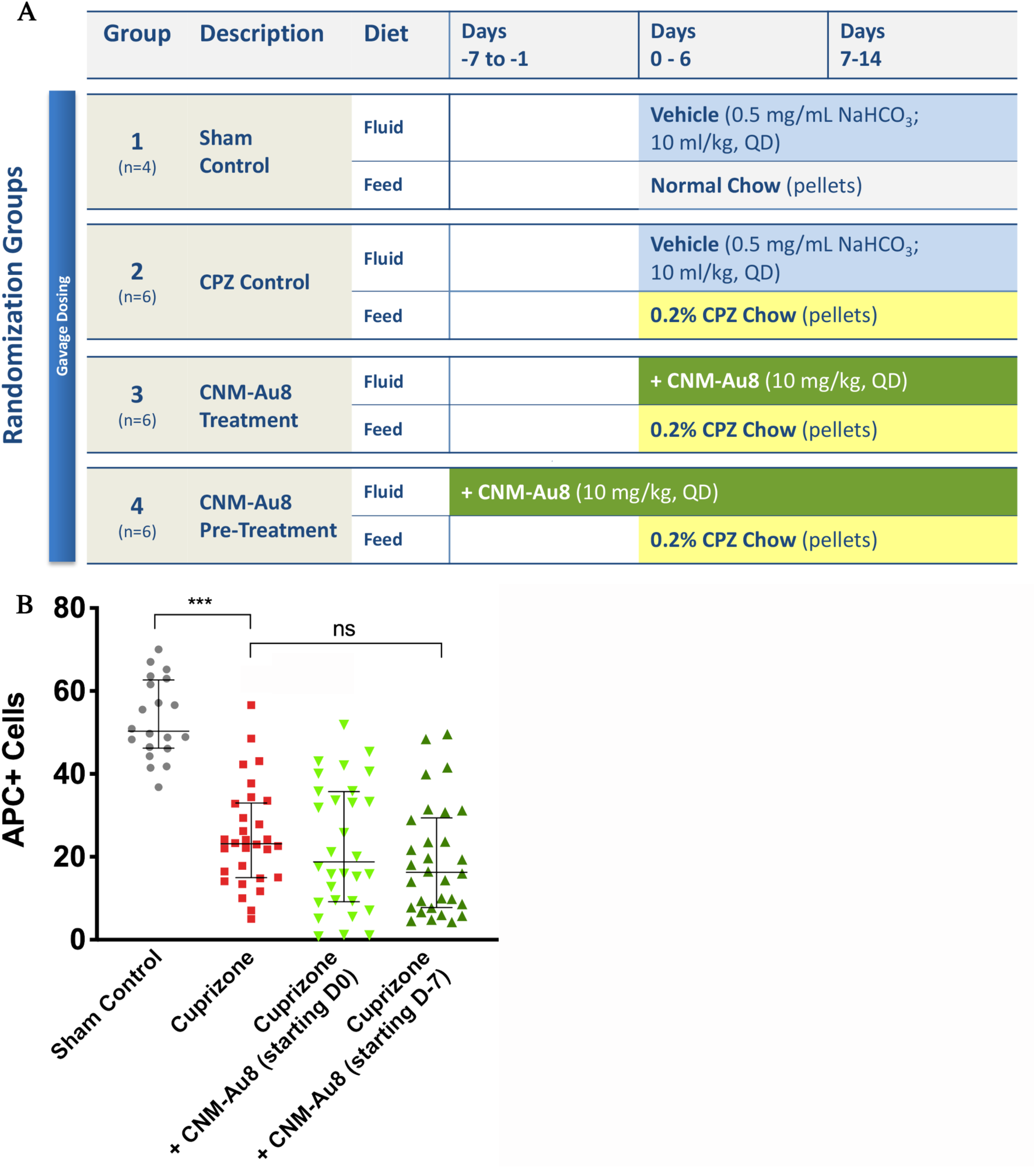
Loss of Apc+ OLs was unaffected by CNM-Au8 treatment within the first two weeks of cuprizone treatment. B, Quantitation of the number of APC positive cells by immunohistochemical staining of coronal sections from each treatment group. Error bars show median and interquartile range.

### Effects on motor functions in cuprizone-treated mice in open-field and movement kinematic behavioral assays

To determine whether CNM-Au8 treatment was capable of restoring function after cuprizone insult, open field gross motor assessments and fine motor kinematic assessments were conducted on groups of mice (N=12 per group, open field; N=10 per group, fine motor kinetics) treated with CNM-Au8 or vehicle. Group 1 was a sham, non-cuprizone control with vehicle delivered to animals by gavage starting on Day 1. Group 2 animals were administered cuprizone and vehicle by gavage for six weeks; Group 3 was administered cuprizone by gavage for six weeks and CNM-Au8 for the final four weeks of the six-week study period. Behaviors for the open field test were assessed prior to cuprizone dosing, at the end of week 3, and at the end of week 6 (Fig 4A), while the fine motor kinematic tests were assessed at the end of week 3, and at the end of week 6.

**Figure 4:**
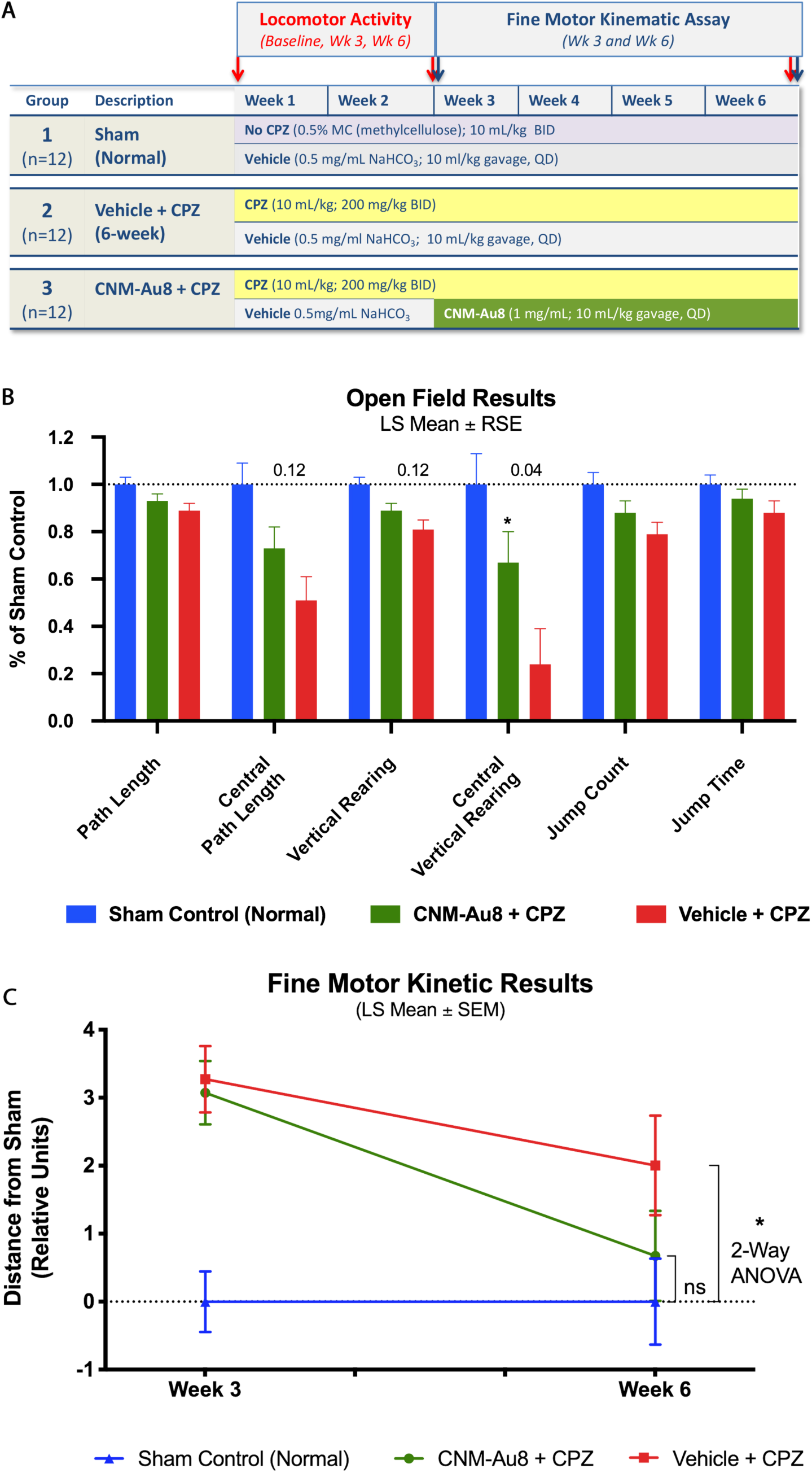
Restoration of function by CNM-Au8 in open field test and fine motor kinematic analyses. A. Quantitation of parameter measures from open field assessments of behavior in untreated sham mice (blue bars), gavage-administered CNM-Au8 treated animals with cuprizone (green bars), and gavage-administered vehicle treated animals with cuprizone (red bars) expressed as a percentage of sham measurements at Week 6. B. Principal component analysis of gait metrics showed no statistical difference (p=0.47) between CNM-Au8 and sham treated groups as compared to a detectable difference in vehicle treated groups vs. sham (p=0.032; 2-way ANOVA) by week 6. See Materials and Methods for statistical calculations.

In eight of the nine quantitative behavioral parameters that were assessed in the open field test, cuprizone-treated animals showed statistically significant behavioral changes from sham-treated animals by the end of six weeks. The ninth metric, average speed, showed insignificant changes between cuprizone-treated and the sham non-cuprizone treated controls at the end of six weeks, and was therefore not evaluated further. Each cluster of bars in Figure 4B shows the change in specific parameter for each group at week 6 compared to baseline as a least-square mean result. Overall, CNM-Au8 treatment of cuprizone-injured animals improved the performance of the mice in each of the open field test assays, substantially restoring the observed cuprizone deficits versus sham treated animals, even though most individual behavioral metrics did not reach statistical significance. More specifically, we observed between 43% to 62% relative recovery toward sham behavior in each parameter due to CNM-Au8 treatment versus vehicle cuprizone treated animals (Table S1), and in all cases there was a trend in improvement noted. Notably, there was a statistically significant improvement in central vertical rearing counts (p < 0.05), and there were near-significant trends in central path length and vertical rearing counts (Fig. 4B).

**Table S1.** Functional improvements in each open field parameter toward Sham controls by CNM-Au8 treated groups compared to vehicle controls. RSE, relative standard error. ^a^p = 0.12. *p=0.04, two-tailed t-test, corrected for multiple comparisons.

|  | **CNM-Au8 plus**<br>**Cuprizone, % of**<br>**sham (RSE)** | **Vehicle plus**<br>**Cuprizone, % of**<br>**sham (RSE)** | **Percent Relative**<br>**Improvement of**<br>**CNM-Au8**<br>**Treatment vs.**<br>**Vehicle**<br>**(Toward Sham)** |
| --- | --- | --- | --- |
| **Path length** | 93.17<br>(-0.03) | 89.02<br>(-0.03) | 62% |
| **Central Path**<br>**Length** | 72.65<br>(0.09) | 51.19<br>(-0.10) | 56%<sup>a</sup> |
| **Vertical Rearing** | 89.03<br>(-0.03) | 80.54<br>(-0.04) | 56%<sup>a</sup> |
| **Central Vertical**<br>**Rearing** | 67.45<br>(-0.13) | 24.41<br>(-0.15) | 43%<sup>*</sup> |
| **Jump Count** | 87.50<br>(-0.05) | 79.23<br>(-0.05) | 60% |
| **Jump Time** | 94.25<br>(-0.04) | 87.87<br>(-0.05) | 47% |

In addition to the open field test, fine motor kinematic analyses were conducted in which parameters measuring limb joint movements, posture, and gait were measured using high-speed cameras that filmed each mouse from multiple viewpoints (top, bottom, and side) as the animal moved. Hundreds of individual measurements from these videos which were then combined to quantitatively describe aspects of gait. Kinematic measurements were made on mice at 3 weeks and 6 weeks after the start of cuprizone treatment. Principal component analysis of these parameters, plotted as distance from sham at weeks 3 and week 6 is shown in Figure 4C. Impairment due to cuprizone can clearly be seen at the end of 6 weeks, with limited overall change vs. sham (-0.67 ± 0.9 relative units) from the week 3 deficit relative to sham values. At Week 6 the cuprizone-treated animals remain impaired when compared to the untreated sham group; -2.0 ± 0.9 units, p=0.032 vs. sham, 2-way ANOVA). In contrast, treatment with CNM-Au8 largely restored kinematic movements to normal levels as the CNM-Au8-treated group became statistically indistinguishable from sham-treated controls at 6 weeks (-0.67 ± 0.9 units; p = 0.468 vs. sham). Between week 3 and week 6, CNM-Au8-treated mice improved by 78% from their week 3 deficit relative to the sham controls.

### Increased OL migration to lesion sites and increased myelin production by nanocrystalline gold following lysolecithin induced demyelination

Lysolecithin injection is an acute, focal method used to induce demyelination. To determine the effect of CNM-Au8 on remyelination in this model, the lysolecithin detergent lysophosphatidylcholine (LPC) was injected into the dorsal white matter of thoracic (T7) spinal cords of rats (N=15 per group). Group 1 was therapeutically treated with CNM-Au8 by gavage three days post-LPC injection and Group 2 was similarly administered vehicle control. Animals were sacrificed for histological or immunohistochemical staining on Day 14 following LPC injection. Histological sections of rat spinal cords at the site of injection were stained with either Luxol Fast Blue (LFB) at day-7 (Fig. 5A-E) or toluidine blue at day-14 (Fig. 5F, G) to visualize myelin. LFB staining revealed the presence of new myelin within the site of the lesion in sections of the CNM-Au8-treated animals (Fig. 5D, E) as compared to the vehicle-treated controls (Fig. 5B, C). Similarly, toluidine blue staining (Fig. 5F, G) shows a striking ‘honeycomb’ pattern of newly myelinated axons within the lesion site, in comparison with a lack of evidence of new myelin in vehicle-treated controls (Fig. 5G). APC (CC1) staining of lysolecithin-induced spinal cord lesions revealed clusters of viable, mature OLs within lesion sites in CNM-Au8-treated animals which were again lacking in vehicle treated controls (Fig. 5H, I).

**Figure 5:**
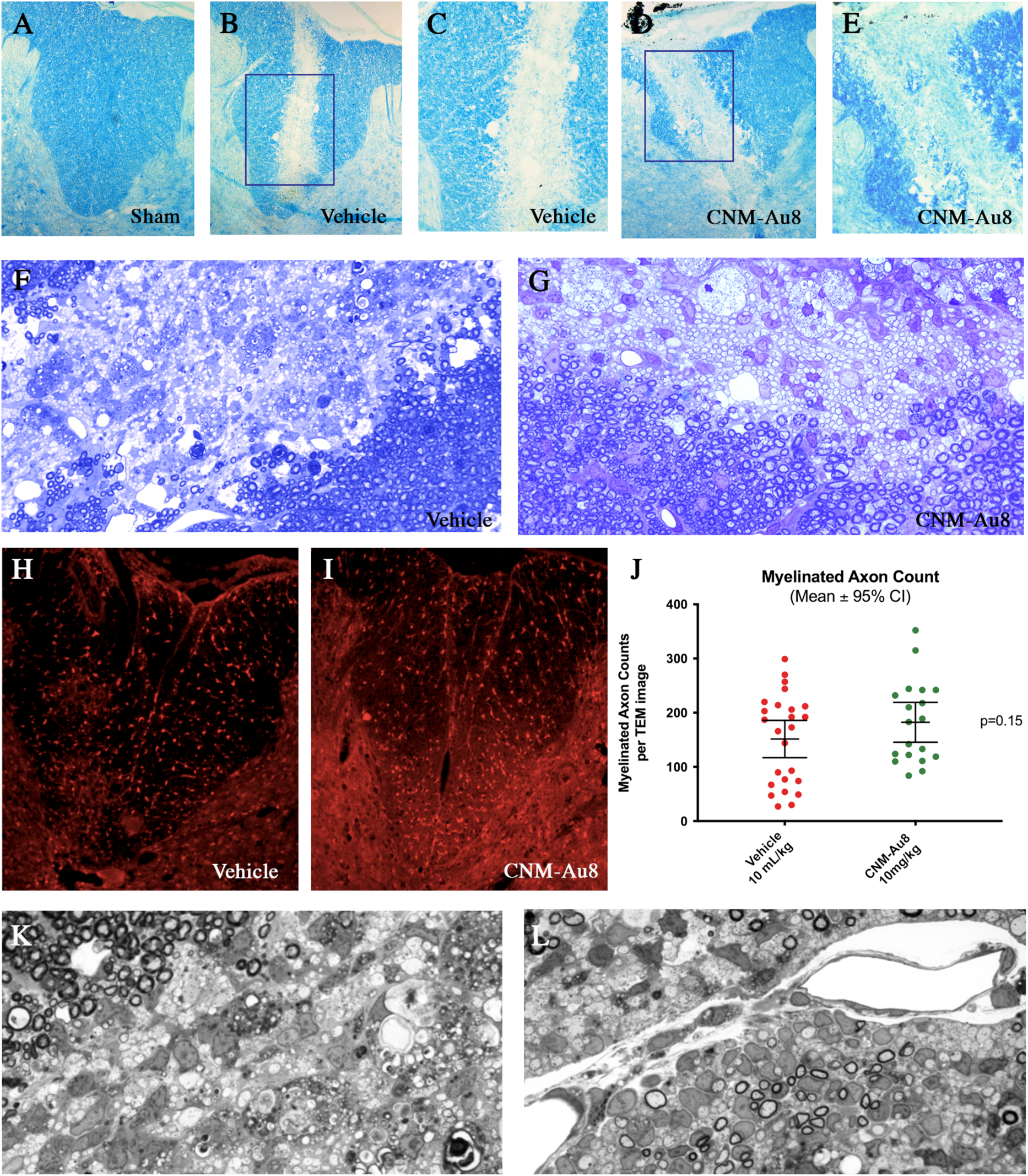
Evidence of remyelination by CNM-Au8 treatment in a focal demyelination lysolecithin rodent model. A-E, Luxol Fast Blue staining of myelin in spinal sections of lesions from sham treated animals (A), lysolecithin lesioned, vehicle treated animals (B and C), and lysolecithin lesioned, CNM-Au8 treated animals (D and E) in which recovery of myelin within lesion area of CNM-Au8 treated animals (D and E) can be seen. A, B, and D: 10x magnification; C and E: 20x magnification. F,G, toluidine blue staining of myelin in spinal sections of lesions from vehicle (F) and CNM-Au8 (G) treated animals showing the distinct ‘honeycomb’ pattern of remyelination in CNM-Au8-treated animals. H, I, anti-APC staining of mature OL cell bodies within spinal sections of lesions from vehicle (H) and CNM-Au8 (I) treated animals. Clusters of mature OLs can be observed in the lesion area of CNM-Au8 treated animals (I). J, Quantitation of myelinated axon counts from TEM images of spinal sections of lesions by animal showed a 43% increase in myelination in treated animals over vehicle controls (p=0.15, unpaired t-test). K,L, Representative transmission electron micrographs of spinal cord lysolecithin lesions of animals treated with vehicle (K) and with CNM-Au8 (L). Normal myelination, at the boundary of the lesion, can be observed in each panel at the upper margins of the field of view. Thinly wrapped remyelinating axons are readily observed in L.

Thinly myelinated axons, which are those with 2-4 wraps of myelin located within lesion sites, indicate axons undergoing remyelination post-LPC injection. A quantitation of thinly myelinated axons within lesion sites was conducted in a blinded fashion using TEM image analyses. The results of the quantitation are shown (Fig. 5J) and indicated that CNM-Au8-treated animals exhibit a 43% mean increase in myelinated axons within lesions post-LPC injection compared to vehicle controls. Representative TEM images clearly show evidence of remyelinating axons within lesion sites of CNM-Au8 treated animals compared to vehicle controls (Fig. 5K, L).

### In vitro differentiation of OPCs with treatment of nanocrystalline gold

We next sought to determine the cellular mechanism underpinning CNM-Au8-mediated remyelination and restoration of motor functions. Our cuprizone and lysolecithin experiments suggested that CNM-Au8 either enhanced the proliferation of OPCs, or enhanced the differentiation of OPCs into OLs, or both. In order to address these possibilities, we utilized an isolated OPC primary cell culture model. OPCs from postnatal mouse cortices were dissociated with papain and isolated using an immunopanning procedure (*21*), then cultured in the presence of CNM-Au8 or vehicle, then counted by flow cytometric methodology following immunohistochemical staining for differentiation markers (*22*).

Determination of the viability of cells in the presence of CNM-Au8 or vehicle was assessed first. OPCs and differentiated OLs were grown in the presence of 0.06, 0.2, 0.6, and 2 µg/mL CNM-Au8 for four days and cell viability was measured using annexin V and propidium iodide stains to differentiate live, apoptotic, and dead cells. No change in viability of OPCs or OLs in response to CNM-Au8 treatment was observed (Fig. 6A, B).

**Figure 6:**
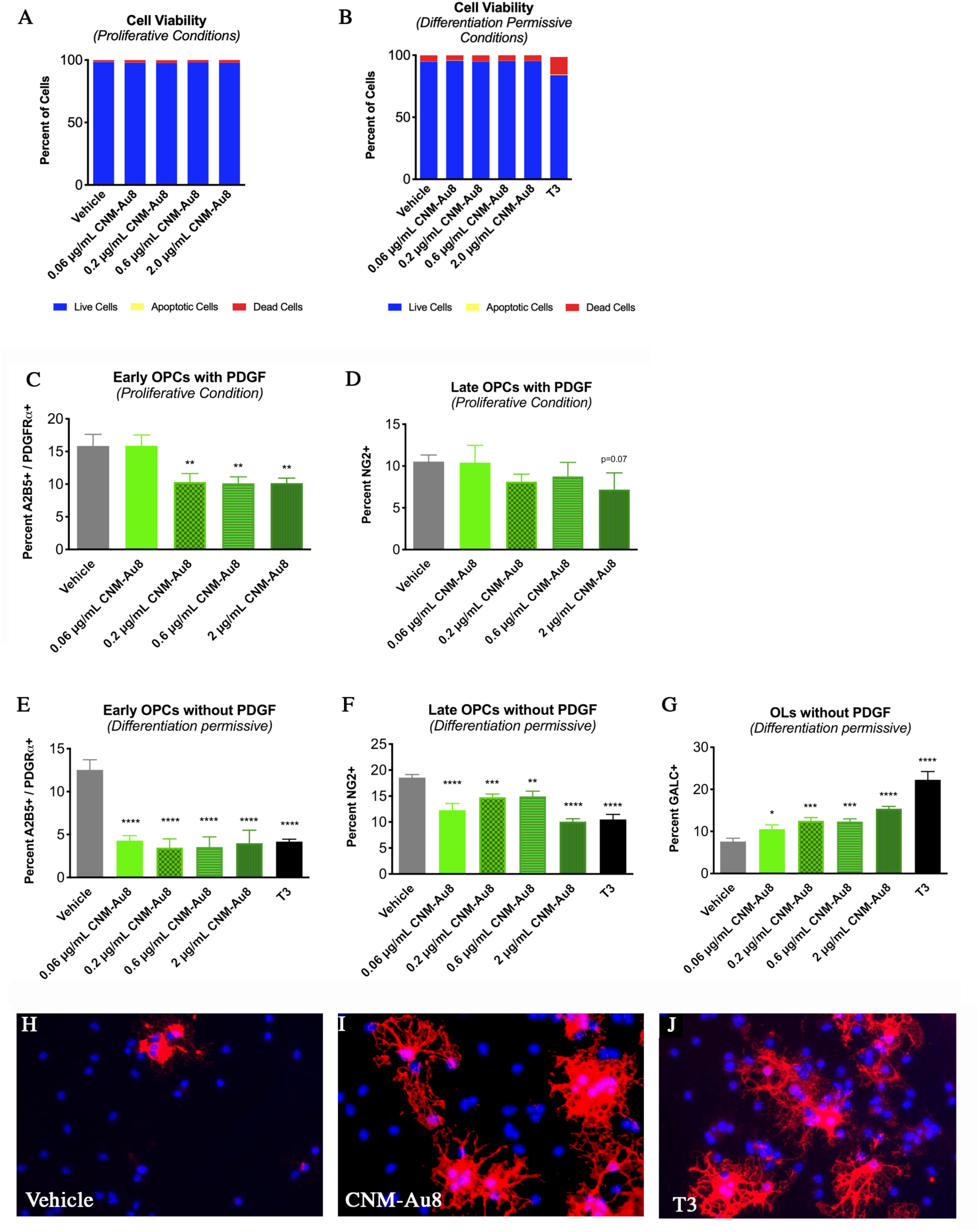
CNM-Au8 mediated differentiation of immunopanned primary OPCs in culture. Increasing concentrations of CNM-Au8 did not affect OPC (A) or OL (B) viability in culture. C and D, OPCs supplemented with 0.01 μg/mL PDGF, ‘proliferative conditions,’ do not show a proliferative response when provided with CNM-Au8 in the media compared to vehicle treated controls; cells expressing markers for early (C) and late (D) OPCs show a relative decline in numbers in response to increasing concentrations of CNM-Au8. E-G, OPCs grown without PDGF, ‘driving conditions,’ show a relative increase in cell numbers expressing the mature OL marker GALC (G), while showing a relative decrease in numbers of OPCs expressing early (E) and late (F) OPC markers compared to vehicle treated controls. H-J, representative images of OPCs treated with vehicle (H), 0.6 μg/mL CNM-Au8 (I) and 0.04 μg/mL T3 (J) stained with DAPI to mark cell nuclei and anti-MBP to mark OLs expressing the mature myelin marker. OPCs treated with CNM-Au8 and cells treated with the positive control T3 showed higher numbers of mature OLs with complex cytoplasmic networks indicative of differentiated OLs.

The proliferative effect of CNM-Au8 treatment on OPCs was assessed by flow cytometry. Results indicated that CNM-Au8, similar to the known OPC differentiation signaling molecule triiodothyronine (T3), suppressed the proliferation of early OPCs (Fig. 6C, D). OPCs were treated for four days with CNM-Au8 in growth media with (Fig. 6C, D) or without (Fig. 6E-G) the proliferation inducer PDGF, stained with antibodies to mark early OPCs (A2B5+/PDGFRα+) or late OPCs (NG2+) and then separated based on marker expression and quantitated by flow cytometry. The percent of OPCs expressing early or late OPC markers was decreased in response to CNM-Au8 or T3 treatment, regardless of whether PDGF was added or not, in comparison with vehicle controls (Fig.6C-G).

In contrast, CNM-Au8 elicited a clear differentiation response from OPCs when cultured in medium without PDGF (Fig. 6E-G). Flow cytometry based on markers for early OPCs (A2B5+/PDGFRα+), late OPCs (NG2+) and mature OLs (GALC+) showed a pronounced shift toward mature OLs in response to CNM-Au8 treatment, as indicated by proportionately higher populations of GALC+ OLs in CNM-Au8-treated cultures (Fig. 6G). Immunohistochemical staining of OLs for MBP further confirmed not only the presence of more MBP+ OLs in response to CNM-Au8, but also that the terminally differentiated cells take on the morphological characteristics of differentiated OLs in the form of extensive networks of processes to a similar extent and degree as the differentiation signaling molecule, T3 (Fig.6H-J).

A second study further confirmed that CNM-Au8 stimulated the differentiation of OLs regardless of whether they were derived from mouse CNS or rat spinal cord cultures. Rat spinal cord cells were dissociated, cultured, and treated with CNM-Au8 or vehicle for 72 hours and then stained with DAPI, as well as antibodies against Olig2 to mark OLs and MBP to mark cell bodies and processes of mature OLs (Fig. S5).

**Figure S5.**
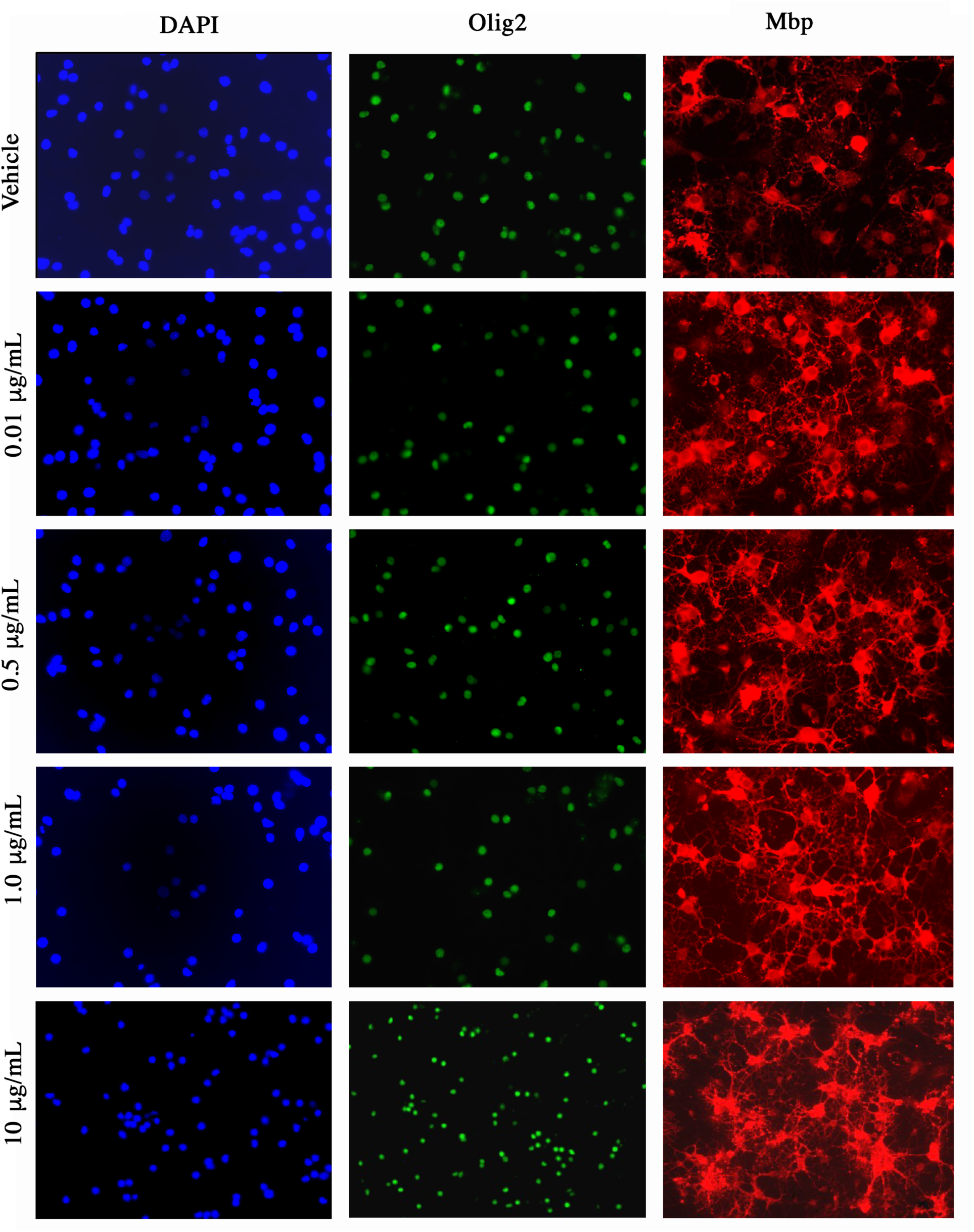
Differentiation of OLs from primary rodent spinal cord cultures by CNM-Au8. Postnatal spinal cord cells were cultured for 72 hours in the presence of vehicle or doses of CNM-Au8 (0.01 μg/mL, 0.5 μg/mL, 1.0 μg/mL. and 10 μg/mL). Cells were then fixed and stained with DAPI, anti-Olig2 antibodies, and anti-Mbp antibodies. Increasing levels of Mbp expression as well as an increase in OL ‘footprint’ can be observed in association with higher levels of CNM-Au8 treatment.

### Differential gene expression analysis of CNM-Au8-treated OPC cultures in vitro

To further characterize the effect of CNM-Au8 on OPC differentiation, we conducted an *in vitro* RNAseq expression study using isolated, purified OPC cultures, with each treatment condition performed in triplicate (Fig. 7). We conducted Multidimensional Scaling Analyses to examine relative distances between samples according to their gene expression profiles. These analyses demonstrated that 1 µg/mL and 10 µg/mL CNM-Au8-treated OPC expression profiles were more similar to the expression profile of differentiated OLs treated with a promoter of OPC differentiation, triiodothyronine (T3), than to the profile of proliferating OPCs treated with PDGF (Fig.7A). Furthermore, CNM-Au8 treatment of mouse OPCs in primary culture for 72 hours results in differential expression (DE) of genes involved in myelination. Markers of oligodendrocyte (OL) maturation such as *Mag*, *Mbp, Gjc2*, *Nkx6.2*, and *Sox10* mRNAs were elevated at least 2-fold over untreated vehicle controls (Fig. 7B). Notably, there was also an enrichment of mRNA transcripts with gene ontologies related to lipid metabolism uniquely present in the DE gene profile for CNM-Au8-treated OPCs (Fig. 7C). These included genes encoding proteins involved in long chain fatty acid synthesis, which is essential to the generation of lipids that comprise ∼70% of myelin. In contrast, the DE gene profiles for the T3 and PDGF controls did not demonstrate enrichment in long chain fatty acid synthesis mRNAs. Taken together, these data demonstrate that treatment with CNM-Au8 promotes OPC differentiation and OL maturation. 

**Figure 7.**
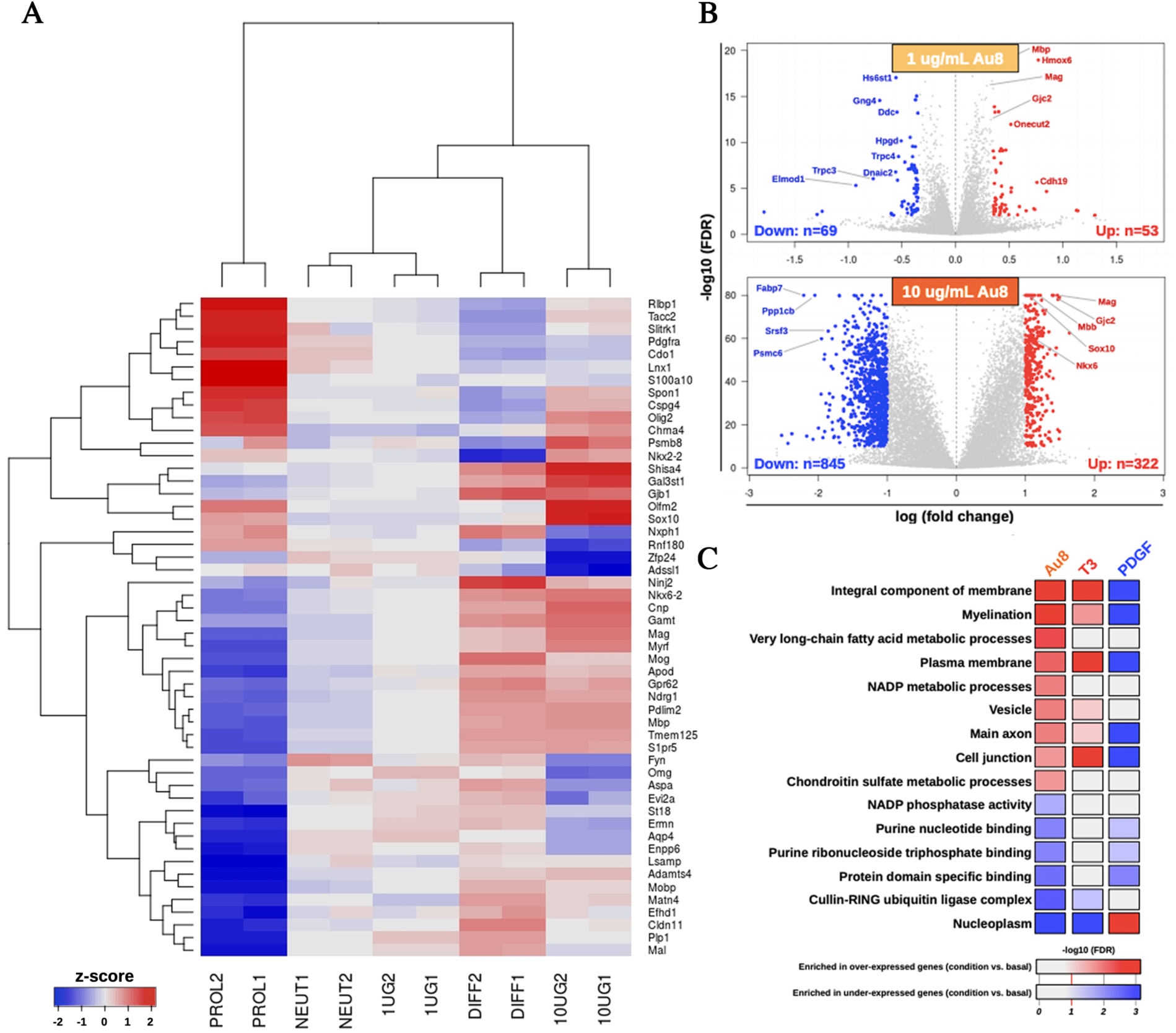
Transcriptomics analyses identified OL differentiation and myelination pathways that are upregulated in response to CNM-Au8. Isolated OPC cultures were treated with vehicle (NEUT1 and NEUT2), 1.0 (1UG1 and 1UG2) or 10.0 μg/mL (10UG1 and 10UG2) CNM-Au8, 0.01 μg/mL PDGF to induce proliferation (PROL1 and PROL2), or 0.04 μg/mL T3 (DIFF1 and DIFF2) to induce differentiation, for 72h in duplicate. Cells were then processed for RNAseq analyses. A, The top fifty variable genes in the dataset are displayed in the heatmap, with the name of each gene in each row labeled to the right. Samples were hierarchically clustered (dendrogram at top), demonstrating that CNM-Au8 treated cells’ transcript profiles are closer, but not overlapping, that of T3-treated cells, as compared to PDGF treated cells. B, Volcano plots of differentially expressed genes, with the identities of upregulated genes known to play a role in differentiation and myelination indicated in red. C, Gene ontology enrichment analysis revealed that gene pathways associated with myelination, plasma membrane regulation, and fatty acid metabolism were among those upregulated upon treatment with CNM-Au8, as well as, in part, with T3.

## Discussion

Gold has traditionally been considered biologically inactive. Ancient cultures took note of its resistance to corrosion, using it to symbolize longevity, and including powdered gold in recipes for life-extending elixirs. Indeed, the stability of gold has been used in a variety of ways for the improvement of human health for over 2500 years, which has presently culminated in the modern-day applications of gold related to bio-imaging, biosensors, and drug delivery (*23*). More recently, advancing technologies for the manufacture of gold nanoparticles have revealed new properties specific to nanoparticulate gold, namely, its catalytic abilities at the nanoscale. In addition to oxidation of NADH to NAD+ first characterized by Huang et al (*12*) and discussed above, gold nanoparticles have demonstrated peroxidase, oxidase, catalase, and super oxide dismutase-like properties, all of which have significant relevance to biologic systems (*24, 25*). We now propose that a novel preparation of clean-surfaced faceted crystalline gold has direct therapeutic ability to support intracellular bioenergetic activities.

We sought to harness gold’s potent catalytic properties at the nanoscale, while minimizing the particles’ potential adverse effects, by devising a novel method of nanocrystal creation using an aqueous electrocrystallization process to grow highly faceted, pure gold crystals that did not rely on any capping or stabilizing agents. This new method afforded several significant advantages: 1) the nanocatalytic activity of CNM-Au8 was significantly enhanced compared to that of standard citrate-reduced gold nanoparticles (Fig. 1C), 2) CNM-Au8 nanocrystals were non-toxic to OPC and OL cells in culture (Fig. 6A, B), and 3) CNM-Au8 functionally improved motor behaviors in mice *in vivo* following exposure to the toxin, cuprizone (Fig. 5). In contrast, gold nanoparticles made using traditional methods with added capping and surfactant agents have been shown to introduce adverse effects in other biological systems (*17*).

Here we have described the creation of a stable CNM-Au8 suspension with highly faceted nanocrystals of an average diameter of 13 nm that are free of surfactants and other modifying chemicals. In several *in vivo* remyelination assays, treatment with CNM-Au8 significantly improved not only the quantifiable detection of myelinated axons in the brains of experimental animals, but also significantly improved mouse behaviors and functional movements in the open field test and kinematic assays. The currently available FDA-approved drugs for the treatment of MS act to suppress the recurrent autoimmune attack on myelin during disease progression. These drugs generally limit disease progression by reducing the frequency and intensity of autoimmune attacks. Used alternately or in conjunction with such drugs, a remyelinating therapy opens up the possibility of restoration of functions that were previously impaired or lost due to MS disease activity, thereby improving patients’ quality of life and potentially reversing disease progression. CNM-Au8 appears to act through an entirely novel mechanism we term ‘nanocatalysis,’ which improves the bioenergetics of nervous system cells through a pathway involving the energetic coenzymes NAD+ and NADH. NAD+ and NADH are vital for driving cellular reduction-oxidation (redox) reactions in living cells in both the glycolytic as well as the oxidative phosphorylation pathways of energy generation. In addition to its redox role, NAD+ also acts as a binding partner and substrate for key regulators of metabolism and repair (*26*). NAD+ is also a precursor for NADP+/NADPH, which are required for cellular anabolic pathways including the synthesis of long chain fatty acids that are required not only for myelin production, but also to protect cells from reactive oxygen species (ROS) (*27*). A NAD+ deficiency of more than 50% has been detected in the serum samples of MS patients compared to healthy controls (*28*). Intriguingly, this NAD+ deficiency increases amongst MS subgroups with increasing severity of the disease (*28*). NAD+ has been proposed to be a potential therapeutic target for MS treatment (*28–30*).

NADH supplementation has been studied as a therapeutic for neurodegenerative diseases in the past, for example for Parkinson’s Disease (PD). However, the promising results of an initial small study using NADH supplementation in 34 PD patients (*31*) were not able to be replicated independently (*32*). A NAD-regenerative component accompanies a catalytic therapeutic such as CNM-Au8, thereby significantly increasing the potential impact on a NAD-deficient system. In addition, use of CNM-Au8 does not preclude additional NAD supplementation; and together, NAD supplementation with CNM-Au8 may act synergistically. Additional studies to explore these possibilities are warranted.

Aerobic glycolysis, even more so than oxidative phosphorylation, plays a key role in oligodendrocyte remyelination (*4*). The reaction provides both energy in the form of ATP as well as pyruvate, a precursor for many myelin-associated molecules (*4*). It has been suggested that the OPCs present in MS lesions are unable to remyelinate because they have undergone cellular stresses that have led to their bioenergetic failure (*3*). We have demonstrated that CNM-Au8 treatment of isolated OPCs in culture stimulated the differentiation of OPCs to mature OLs. This activity can explain the enhanced number of mature OLs we observed in demyelinated lesion sites in response to CNM-Au8 treatment in our animal models.

Many neurodegenerative diseases, including Alzheimer’s, Parkinson’s, and Huntington’s diseases are all thought to involve bioenergetic failure, which manifests as the inability to efficiently protect the brain from excitotoxic events associated with neural activity, inability to modulate reactive oxygen species, and inability to continuously provide adequate energy stores for normal neuronal function as aging progresses (*33*). Moreover, a ^31^P magnetic resonance spectroscopy study has shown that NAD+/NADH redox potential declines in the aging brain, and lowered NAD+/NADH redox potential has been observed in patients with schizophrenia compared to age-matched, gender matched controls (*34, 35*).

There are other remyelinating agents in various stages of clinical development, including opicinumab, an anti-LINGO monoclonal antibody; elezanumab, an anti-RGMa monoclonal antibody; NDC-1308, an estradiol derivative; and clemastine fumarate, an older generation antihistamine and M1/M3 muscarinic receptor reverse antagonist (*36–38*). Remyelination is a complex process involving multiple intrinsic and extrinsic signals to the OL (*37*). Accordingly, these therapies work by blocking inhibitory signaling pathways, or by stimulating positive signaling pathways, of OPC differentiation (*38–40*). Unlike these signaling therapies, CNM-Au8 catalytically compensates for energy loss, stimulating OPC differentiation and accelerating myelination in a unique mode of action that likely falls downstream of many stimulatory signaling pathways (*29, 30*). Moreover, bioenergetic failure is a noted pathophysiologic mechanism common to many age-related neurodegenerative diseases (*33*). Because CNM-Au8 does not target one specific pathway or cell type in the CNS, the applicability CNM-Au8 as a neurotherapeutic is potentially wide-ranging and highly significant to many neurodegenerative diseases impacted by impaired cellular bioenergetics.

There are limitations to this study. While we demonstrated an improvement in open field and fine motor behaviors in response to CNM-Au8 in cuprizone-treated mice, many of the individual parameters measured did not reach statistical significance distinct from the cuprizone-treated controls (Fig. 4). Post-hoc sample size calculations based on the observed differences between the Group 1 (sham) animals and the Group 2 (vehicle plus cuprizone treated) animals indicated that significantly more animals would be needed for an appropriately powered experiment. Indeed, the inherent variability of the effects of cuprizone on the corpus callosum indicated that even in the large (N=15 mice per group) remyelination replication study (Fig. 2) a higher powered study would better be able to statistically distinguish morphometric differences in remyelinated axons in the CNM-Au8 treated groups versus vehicle. At this point in time, we do not have a clear explanation as to why the therapeutic arm treatment groups outperformed corresponding groups in the prophylactic arm, in spite of the fact that animals in the prophylactic arm were treated with CNM-Au8 for a longer duration. Cuprizone itself does not consistently demyelinate brain areas uniformly when evaluated both on a within-animal and a between-animal basis (*41*); therefore the toxin’s effects, particularly if mild, can be difficult to demonstrate on a quantitative basis. Nevertheless, remyelination due to CNM-Au8 treatment was reproducibly demonstrated in all five cuprizone and the lysolecithin models presented herein, when evaluated on ultrastructural, cellular, and functional levels. The observed functional improvements with oral administration of CNM-Au8 in every parameter measured in the open field test, and the overall gait improvement observed using fine motor kinetics are encouraging indicators of efficacy that future studies will be able to confirm.

While we demonstrated the remyelinating capabilities of CNM-Au8 treatment using two independent models in mice and rats, namely cuprizone and lysolecithin, a third widely-used demyelinating model of experimental autoimmune encephalomyelitis (EAE) model produced inconsistent results with CNM-Au8. In contrast to the cuprizone and lysolecithin models, the EAE model involves the induction of an autoimmune attack on the animals’ myelin by injection of myelin proteins subcutaneously, resulting in an aggressive, continual autoimmune attack with no periodic times for remyelination recovery. The very nature of this experimental model may be the reason for our experiments using the EAE model being inconclusive, since the extensive axonal loss associated with chronic EAE and subsequent neurological deficit may preclude the possibility of a remyelinating therapy demonstrating efficacy in these models due to the lack of sufficient substrate (axons) for which OLs can remyelinate (*42, 43*). Some have raised concerns that the model does not accurately reproduce key features of MS pathology; and even though EAE models led to the development of some MS therapies such as glatiramer acetate and natalizumab, numerous other candidates that showed promise in EAE models were found to be detrimental or ineffective for MS (*44*).

In addition to the preclinical efficacy studies of CNM-Au8 as a potential remyelinating therapeutic for MS described here, the drug development program for CNM-Au8 included IND-enabling toxicology studies as well as a First-in-Human Phase I study. The results of the genotoxicity, safety pharmacology, and acute and chronic toxicology studies for six months’ duration in rodents and nine months’ duration in canines indicated a clean safety profile for CNM-Au8, with no severe adverse effects identified in any of these studies even at the highest maximum feasible doses tested. A Phase I First-In-Human study of CNM-Au8 was completed in October 2016 (Study AU8.1000-14-01; NCT02755870), involving 86 healthy volunteers and two study arms, a single ascending dose and multiple ascending dose arms. All dosing regimens were well-tolerated in this study, with treatment-emergent adverse events reported as predominantly mild. There were no serious adverse events or events leading to treatment discontinuation. Due to promising preclinical data described here and the favorable toxicity profiles achieved by CNM-Au8, a Phase II, double-blinded, randomized, placebo-controlled VISIONARY-MS (Treatment of Visual Pathway Deficits In Chronic Optic Neuropathy for Assessment of Remyelination in Stable Relapsing-Remitting Multiple Sclerosis) is currently underway.

## Materials and Methods

### Animal care and husbandry

All procedures were performed with the approval of the Hooke Labs, Charles River Labs, Northwestern University, and the George Washington University Institutional Animal Care and Use Committees, and all laboratories adhered to the NIH Guide for the Care and Use of Laboratory Animals in accordance with Federal Animal Welfare Regulations. All animals were housed under pathogen–free conditions with food and water provided *ad libitum,* except as noted in text when CNM-Au8 or vehicle was provided as animals’ sole fluid source instead of water. Gavage administered animals were still provided water *ad libitum*. C57BL/6 was the mouse strain used in all experiments involving mice or primary cell cultures from mice. Sprague-Dawley rats were used in all experiments involving rats or primary cell cultures from rats.

### Vehicle

Vehicle used in all experiments is 6.5 mM NaHCO_3_. In gavage experiments, volume of vehicle delivered was always equivalent to the gavage dose of CNM-Au8 (10 mL/kg).

### Cell-free NADH oxidation assays

Oxidation of NADH to NAD+ in presence of CNM-Au8 was monitored by UV-VIS absorbance spectra as previously described (*12*). NADH disodium salt (bioPLUS, Dublin, OH) was dissolved in DI water to a concentration of 1mM stock solution. NIST 30nm and NIST 10 nm were acquired from the National Institute of Standards and Technology (Gaithersburg, MD). UV-VIS Spectra were exported into MS Excel and analyzed with Origin 6.0 software.

### In vitro quantitation of NAD+ and NADH levels

Rat mesencephalic neurons and glia were cultured as described (*45*). Briefly, midbrains obtained from 15-day-old rat embryos were dissected under a microscope. Midbrains were removed and dissociated by trypsinization for 20 min at 37 C. The reaction was stopped by the addition of Dulbecco’s modified Eagle’s medium (DMEM) containing DNAse I grade II (0.5 mg/mL) and 10% fetal calf serum (FCS). Cells were mechanically dissociated by three passages through a 10 mL pipette, then centrifuged at 180 x g for 10 min at +4 C on a layer of BSA (3.5%) in L15 medium. The supernatant was discarded and cell pellets were resuspended in Neurobasal (Invitrogen) supplemented with B27 (2%), L-glutamine (2mM), 2% of P/S, 10 ng/mL brain-derived neurotrophic factor (BDNF), and 1 ng/mL glial-derived neurotrophic factor (GDNF). Viable cells were counted in a Neubauer cytometer using the trypan blue exclusion test. Cells were seeded at a density of 40,000 cells/well in 96-well plates pre-coated with poly-L-lysine and maintained in a humified incubator at 37 C in 5% CO_2_. After four days of culture, CNM-Au8 (10ng/mL, 100 ng/mL, 1 μg/mL, 10 μg/mL final concentrations), BDNF (50 ng/mL), or vehicle was added for 36 h. Quantitation of NAD+ and NADH was performed using Promega’s NAD+/NADH-Glo quantification kit (cat # G9071).

### In vitro quantitation of ATP levels

The human oligodendrocytic hybrid cell line M03.13 (TebuBio, 007CLU301-A) was grown in low glucose DMEM (5mM glucose, PAN BIOTECH P04-01550) with HEPES 10mM and 10% FBS. Cells were seeded at a density of 20,000 cells per well in a 96-well plate, and then treated with CNM-Au8 (0.1 µg/mL, 0.32 µg/mL, 1 µg/mL) or vehicle for 72h. Prior to lysing cells, half of the wells (N=3 per condition) were treated with a mitochondrial blocking cocktail (100 µM antimycin A, 0.5µM rotenone, 3 µM oligomycin) for two hours, then immediately lysed in buffer containing luciferase. ATP concentration in lysates was determined by measurement of bioluminescence using a 1s integration time on a luminometer (CLARIOstar, BMG LABTECH). Mitochondrial ATP production was calculated by subtracting the average ATP level from mitochondria-blocked cells from average total ATP levels measured in untreated cells.

### Extracellular Acidification Rate (ECAR) measurements

Murine OPCs derived from P7 mouse pups were immunopanned as described above and seeded in 96-well Agilent plates in OPC optimal medium (*21*) for one day to allow for stabilization. CNM-Au8 (0.3 ng/mL, 1 ng/mL, 3 ng/mL, 10 ng/mL) or vehicle was added to the culture for 24 h. Aerobic glycolysis, as determined by extracellular acidification rate (ECAR) of culture media, was measured using the Seahorse flux analyzer (Agilent). Protein content was determined using the Bradford assay (BioRad). Experiments were performed in triplicate.

### Cuprizone treatment

The 5-week cuprizone demyelination model was used as described (*20*). One hundred and five male mice (Taconic Farms, 8 weeks old) were assigned to one of seven groups, with each group balanced by weight, N = 15 animals in each group. The 0.2% cuprizone chow (catalog #TD.140803) and normal control chow were supplied by Harlan Laboratories (Bethesda, MD). Groups 1-5 were given vehicle (0.5 mg/mL 6.5 mM NaHCO_3_ at 10 mL/kg by gavage daily) or CNM-Au8 (10 mg/kg/day) by gavage at the same time (+/-1 hour) every day. Cuprizone treatment started on Day 0 for all groups except Group 1, which was fed normal chow for the duration as a negative control. Prophylactic dosing was started on Day 0 for Groups 1-4 and Group 6 and continued to the end of the study. Therapeutic dosing started on Day 14 for Groups 5 and 7 and continued to the end of the study. Group 2 mice were sacrificed on Day 14; all other groups were sacrificed on Days 34-36 (five weeks of treatment). Three mice, one from each of Groups 1, 5, and 7 died before their scheduled termination.

For the post-cuprizone CNM-Au8 treatment study, twenty-five 8-week old, male mice were divided by weight into five groups. For treatment schematic for each group, see Fig. 2A. Groups 2 and 3 were fed 0.2% cuprizone for five weeks, then the feed was replaced with normal chow, and CNM-Au8 (10mg/kg/day by gavage once daily) or vehicle (0.5 mg/mL NaHCO_3_ at 10 mL/kg by gavage once daily) was provided for one (Groups 2a and 2b) or two weeks (Groups 3a and 3b) before animals were sacrificed for immunohistochemical analyses. Group 1 was fed normal chow for the first five weeks and was then provided vehicle by gavage.

### Corpus callosum sectioning and TEM imaging

Mice were anesthetized by Avertin (250-400 mg/kg) injection into the peritoneum then perfused with 50 ml cold saline (0.9%NaCl in dH_2_O) until the liver became completely clear. After fixing tissue with 150 -180 ml of fixative (3.5% paraformaldehyde, 1.5% glutaraldehyde in 0.1M cacodylate), two 2 mm sagittal slices of brain tissue on either side of the midline containing the posterior corpus callosum were dissected. Samples were post-fixed in 1% osmium tetroxide in cacodylate buffer, followed by post-fixation in 1% uranyl acetate (Ted Pella #19481). After stepwise dehydration in increasing concentrations of ethanol, the samples were embedded in resin (Eponate 12; Ted Pella #18005 for sectioning. 90 nm thick sections were cut between Bregma 0.82mm and -1.82 mm using an Ultramicrotome (Reichert Jung Ultracut). Imaging was carried out at 4000x, 16,000x, and 40,000x magnifications using Zeiss Libra 120 transmission electron microscope.

### Coronal brain sectioning, fixing, immunohistochemical staining and quantitation of signal

Mice were deeply anesthetized and transcardially perfused with 30 ml cold PBS followed by 30 ml 4% paraformaldehyde in PBS. Intact brains were removed and post-fixed in 4% paraformaldehyde in PBS overnight. Brains were then cryopreserved in 30% sucrose in PBS approximately 2 days until they sank. A coronal block encompassing approximately 1 mm to either side of bregma was cut, immersed in OCT, then snap frozen. 10 µm coronal sections were cut by cryostat and mounted onto glass slides. Matched coronal brain sections including the body of the corpus callosum (approximately 10 sections per animal) were identified, blocked with donkey serum, and stained for MBP (Abcam, ab40390) or APC (Abcam, ab15270). Sections were stained with 5 µg/ml MBP or 1 µg/ml APC overnight at 4°C, washed in PBS, then stained with AlexFluor 488 or AlexaFluor 594 (Millipore, AP182B, 1:500) for one hour at room temperature. Sections were counterstained with Hoechst. Slides were imaged on a Nikon A1R+ Confocal Laser Microscope and images encompassing the midline of the corpus callosum were acquired. Post-image analysis was conducted using ImageJ software. Briefly, total number of CC1+ cell bodies (at least 8 sections per animal) were counted in a standardized region of interest (ROI) and compared between experimental groups. Density of MBP staining in a standardized ROI of the corpus callosum (at least 8 sections per animal) was determined by computing the integrated density of the fluorescence signal and compared between experimental groups.

### Qualitative assessment of TEM images

Qualitative ultrastructural analyses of over 3000 TEM images (at magnifications of 4000x, 13,500x, 16,000x, and 40,000x) from 85 mice (10-15 animals per treatment group) were carried out by an expert pathologist (S.K.) who was blinded to the treatments of the experimental groups. The data for each animal was presented as a series of images (8 minimum) arranged from lowest to highest power.

### Quantitation of percent myelination and g-ratios from TEM images

Three investigators (B.F, J.F., N.C.) who were blinded to the treatment protocol independently reviewed all available TEMs taken at 16,000x magnification from the corpus callosum of animals from all seven groups [N=587 images total; 70 images from Group 1 (N=7 animals), 110 from Group 2 (N= 15 animals), 79 from Group 3 (N= 8 animals), 94 from Group 4 (N= 9 animals), 90 from Group 5 (N= 9 animals), 53 from Group 6 (N=7 animals), and 91 from Group 7 (N= 7 animals).]. ImageJ was used to count the number of myelinated and unmyelinated axons in each image to calculate the percentage of myelinated axons per image. Because of the high variability of the number of axons in each image, there was high variability in the extent of demyelination due to cuprizone per image as well. Therefore to be able to compare images with similar axon densities against each other, the dataset for each group was divided into quartiles based on the number of myelinated axons of Group 1 control images. G-ratios are calculated as the inner myelin circumference divided by the outer myelin circumference.

### Lysolecithin injection demyelination model

The demyelinating lysolecithin lesion model was used as described (*46*). In brief, local laminectomy exposed the area of T7 spinal cord of deeply anesthetized rats, and 1.5ml of 2% LPC is delivered locally using a pulled glass needle over a 5-min period. Animals were allowed to recover for 7 or 14 days prior to analysis. Vehicle (N=15 rats) or CNM-Au8 (N=15) was administered by gavage daily (10 mg/kg/day) starting on day 3 following lysolecithin injection. Animals were sacrificed on day 7 or day 14 and lesioned areas, marked by charcoal, were sectioned for light microscopy. Sections were stained for myelin using Luxol fast blue or toluidine blue. Quantitation of myelinated axon counts was conducted on TEM images of lesion sections by counting the number of thinly-wrapped (2-4 wraps) axons within lesion boundaries, in blinded fashion.

### Open field and fine motor kinematic studies

Open field (N=12/group) and fine motor kinematic (N=10/group) analyses of cuprizone/CNM-Au8 treated animals were performed by Charles River Laboratories.

For the open field test: Activity chambers (Med Associates Inc, St. Albans, VT; 27 x 27 x 20.3 cm) were equipped with infrared beams. Female mice were placed in the center of the chamber and their behavior was recorded for 30 min in 5-min bins. Quantitative analysis was performed on the following dependent measures: total path length, path length in the center of the open field, total vertical rearing, vertical rearing in the center, total number of jumps, number of jumps in the center, total time spent jumping, time spent jumping in center, and speed.

For fine motor kinematic analysis: Fine motor skills and gait parameters were assessed using a high precision kinematic analysis method (MotoRater, TSE Systems, Homberg, Germany), allowing for automated analysis of general gait patterns, body posture, balance and fine motor skills. Before each test session, mice were externally marked on joints of limbs and tail. The movement data were captured using a high speed (300 fps) cameras positioned above, below, and at the side of the animal. SimiMotion software was used to track the marked points of the body in relation to the ground and in each of the three dimensions. Gait patterns and movements were analyzed using a custom automated analysis system (Charles River, Inc.). The analyzed parameters included general gait pattern parameters (stride time and speed, step width, stance and swing time during a stride, interlimb coordination), 2) body posture and balance (toe clearance, iliac crest and hip height, hind limb protraction and retraction, tail position and movement), and 3) fine motor skills (swing speed during a stride, jerk metric during swing phase, angle ranges and deviations of different joints, vertical and horizontal head movement). The analysis provided 83 different parameters related to fine motor capabilities and gait. Data were analyzed for distinctive individual parameters, as well as by principal component analysis (see Statistical methods).

### Viability assay and flow cytometry proliferation/differentiation assays of immunopanned OPCs

Oligodendrocyte precursor cells (OPCs) from stage P5-7 C57BL/6 mouse pups were immunopanned as described (*21*) in media supplemented with 0.01 μg/mL PDGF (‘proliferative conditions’) or 0.04 μg/mL triiodothyronine (T3) (‘driving conditions’), and treated with vehicle or CNM-Au8 for 72 hours. Flow cytometry was performed as described (*22*).

### RNASeq Transcriptomics

Oligodendrocyte precursor cells (OPCs) from stage P10 C57BL/6 mouse pups were immunopanned and cultured as described (*21*), then treated with vehicle, 0.04 μg/mL triiodothyronine (T3), 0.01 μg/mL PDGF, or 1 ug/mL or 10 ug/mL CNM-Au8 for 72 hours. Two technical replicates were conducted for each condition.

RNAseq libraries were prepared using RNA extracted from treated OPCs then sequenced (Illumina Hiseq4000). Reads were aligned using STAR; gene counts were computed by *htseq-count*. Downstream analyses were performed using R, and the *limma*, *edgeR*, *TopGO* Bioconductor packages. Heatmaps were plotted using the *gplots* R package.

### Statistical methods

Unless described below, all statistical analyses indicated in the text were carried out using statistical calculators included in the standard MS Excel package and/or Prism 7.0. Statistical significance was defined as p values of 0.05 or less.

#### Myelinated-Percent of Normalized Distributions per TEM Image by Quartile

Group 2 (early sacrifice) results were excluded from analysis. The remaining 6 treatment groups used were analyzed independently by quartile. Quartile assignment for each image was based on the distribution per Normal Myelin Axon Count from Group 1. For each treatment group, summary statistics (mean, standard deviation, median, minimum, maximum) of the Myelinated-Percent of Normal Distribution per TEM Image were calculated. To test for treatment differences, analyses using a nested mixed model were carried out. A nested mixed model was used to account for within subject (mice) correlations between images. For the model, the unit of analysis (dependent variable) was the Myelinated-Percent of Normal Distribution per TEM Image. The fixed effect for treatment was included in the model. A random effect was included in the model for the individual mice. In addition, the effect for mouse was nested within treatment group. The Kenward-Roger method was used to compute the degrees of freedom for the test of fixed effects.

Least square (LS) means were presented for each treatment. LS means were calculated to account for the unbalanced data of the model effects. For each LS mean calculated, an associated 95% confidence interval (CI) was presented. Comparisons of treatment groups were carried out by calculating the difference of the LS means. All possible pairwise comparisons that involved Group 1 (no cuprizone, vehicle control) and Group 3 (plus cuprizone, vehaicle control) were carried out. A p-value < 0.05 was taken to be significant.

#### Open Field Parameters Analysis

Endpoints analyzed were the following: Path Length, Central Path Length, Vert. Rearing, Central Vert Rearing, Jump Counts, Jump Time. Each endpoint had results for 3 time points: Baseline, Week 3, Week 6. For each post baseline time point (Week 3, Week 6), change from baseline was calculated. Summary statistics (n, mean, standard deviation, median, minimum, maximum) for both the raw scores and the change from baselines were calculated.

To test for treatment differences (change from baseline), mixed model repeated measures (MMRM) analyses were carried out to account for the longitudinal nature of the data. Covariates included the baseline score of the endpoint of interest and the baseline weight. An unstructured covariance model was used to account for correlated measures within a subject. The Kenward-Roger method was used to compute the degrees of freedom for the test of fixed effects. LS means and an associated 95% confidence interval (CI) for each treatment and time point were calculated to account for the unbalanced data of the main effects and covariates. Comparisons of treatment groups were carried out by calculating the difference of the LS means. A p-value ≤ 0.05 was taken to be significant.

#### Mouse Motor Kinematics

Ten principal components (PC) of gait features were identified from over 100 kinematic gait parameters collected on mice (Charles River Labs – look up ref). Those 10 components were labeled as follows: PC1- Slowness, PC2- Tail Tip Height, PC3- Knee Function, PC4- Step Length, PC5- Interlimb Coordination, PC6- Vertical Movement, PC7- Hip Angle, PC8- Hindlimb Trajectory, PC9- Hip Height, PC10- Forelimb Trajectory. Each principal component had results for 2 timepoints: Week 3, Week 6.

The three treatment groups used for this analysis were Sham, Vehicle, and CNM-Au8. For each treatment group and time point, summary statistics (n, mean, standard deviation, median, minimum, maximum) for the raw scores were calculated.

To test for treatment differences, mixed model repeated measures (MMRM) analyses were used to account for the longitudinal nature of the data. For the model, the unit of analysis (dependent variable) was the principal component. Each principal component was analyzed separately. Fixed effects for treatment and visit and the interaction term treatment and visit were included in the model. A random effect was included in the model for the individual mice. An unstructured covariance structure was used to account for measures within a subject being correlated. The model converged using the unstructured covariance structure. The Kenward-Roger method was used to compute the degrees of freedom for the test of fixed effects.

Least square (LS) means were presented for each treatment and time point. LS means were calculated to account for the unbalanced data of the main effects. For each LS mean calculated, an associated 95% confidence interval (CI) was presented. Comparisons of treatment groups were carried out by calculating the difference of the LS means. A p-value ≤ 0.05 was taken to be significant.

## Supplemental Materials and Methods

### Preparation of CNM-Au8 and characterization of CNM-Au8 (Table 1 and TEM)

Preparation and characterization of gold nanocrystals CNM-Au8 were as described in U. S. Patent 9,603,870 B2.

### Cuprizone experiments

Cuprizone experiments described in the Supplemental section were performed as described in the main paper Materials and Methods, with the following changes:

For the pilot cuprizone experiment, sixteen mice were divided into four groups with N=4 per group. Group 1 served as a negative control with vehicle provided *ad libitum* in drinking water and normal chow. Group 2 was fed 0.2% cuprizone chow with vehicle provided *ad libitum* in drinking water. Group 3 received CNM-Au8 (50 µg/mL) in their drinking water, and normal chow. Group 4 received CNM-Au8 (50µg/mL) in their drinking water, and 0.2% cuprizone chow. After five weeks of treatment, animals were prepared for TEM imaging as described. The average intake of CNM-Au8 in the treatment groups was measured on a daily, per cage basis. From this data, a daily dose of 10 mg CNM-Au8/kg/day was calculated to be a sufficient dose for detectable effects on myelin.

For the early sacrifice cuprizone experiment, four groups of mice were treated as follows: Group 1 (N=4) was given normal chow and vehicle (10mL/kg) by gavage; Group 2 (N=6) was given 0.2% cuprizone chow and vehicle by gavage; Group 3 (N=6) was given 0.2% cuprizone chow and CNM-Au8 by gavage (10 mg/kg); Group 4 (N=6) was given 0.2% cuprizone chow on Day 0, and CNM-Au8 by gavage (10 mg/kg) starting seven days prior to Day 0 when all other groups were started on both cuprizone and CNM-Au8 or vehicle, and continuing through to Day 14. All animals were sacrificed on Day 14 and coronal brain slices were prepared for immunohistochemical staining and quantitation as described in Materials and Methods.

### PLP staining and quantitation

7 µm thick serial coronal brain sections between bregma-0.82 mm and bregma-1.82 mm were prepared and analyzed.

According to the methods referenced previously^3^, paraffin-embedded sections were de-waxed, rehydrated, while housed in a glass container partially filled with 10mM citrate buffer (pH 6.0), and then microwave-heated in a conventional 1.65 KW household microwave until the buffer began to boil. Brain sections were then quenched with 0.3% H_2_O_2_, blocked for 1 hr in PBS containing 3% normal horse serum and 0.1% Triton X-100. Sections were then incubated at 4°C overnight with anti-PLP antibodies (AbD Serotec) at a dilution factor of 1:500. After washing with washing buffer (PBS buffer, pH 7.4), coronal brain sections were further incubated with biotinylated anti-mouse IgG secondary antibody (Vector Laboratories) for 1 hr, followed by exposure to peroxidase-coupled avidin-biotin complex (ABC Kit, Vector Laboratories) for 30min and developed with diamino-3,3’benzidine reagent according to the manufacturer’s instructions (Vector Laboratories). To determine the total cross-sectional area of all brain matter present on each slide, sections were also stained with hematoxylin to label all brain tissue. Specifically, to quantify the amount of immunopositive PLP in the coronal portion of each mouse brain, coronal sections (i.e., between bregma-0.82mm and 1.82mm) were scanned using a specially adapted Cannon Scanner (output resolution of 2400 dpi). Each pixel in the scan was then evaluated and automatically by Photoshop, assigned a value between 1 and 255, with “255” corresponding to the least intensity and “1” corresponding to the highest intensity. The data were then exported to Excel. The total number of pixels assigned each number between 1 and 255 were then tabulated to achieve a histogram. The data for each histogram was then analyzed and a quantitative weighted average for the amount of pixel intensity in each myelin-stained coronal slide was determined. This quantitative number (appropriately corrected for background) resulted in the ability to make a direct comparison of the amount of color for an equal area in each myelin-stained coronal slide.

Further, in order to make a meaningful scientific comparison of the amount of demyelination/remyelination (i.e., color or shade intensity) in each mouse Group, it was also necessary to determine the total amount (i.e., cross-sectional area) of brain matter present in each of the coronal sections. Thus, the total brain matter area on each coronal slide needed to be determined so that the total amount of color/brain matter area could be quantified and normalized. The amount of color per unit area was then used to compare directly the relative amount of myelin present. Accordingly, hematoxylin was used to stain all of the brain matter in another immediately adjacent serial coronal section. These brain sections were similarly quantified and the total “amount of color” (i.e., lightness or darkness) determined in a first coronal slide, was compared to the total cross-sectional area of brain matter present (represented by both blue coloring and yellow-brown coloring intensity above a minimum threshold amount), determined in a juxtaposed or serial coronal slide to determine the total amount of color or shade/unit area of brain matter.

As stated above, there were 8 groups of mice, and, for staining purposes, each group had 3 mice. Because staining intensity varied as a function of time (e.g., when the staining steps were performed) care was taken to normalize staining or color intensity variations.

### CNM-Au8 and cuprizone chelation of copper

Materials: cuprizone (Sigma-Aldrich 14960), copper (II) sulfate, 5-hydrate - Macron (4844-04), copper (II) acetate mono-hydrate (Alfa Aesar A16203), potassium gold(III) chloride (Sigma-Aldrich 334545). 1.0 mM cuprizone solutions were prepared by dissolving solid cuprizone in 50% ethanol. Copper(II) sulfate and copper(II) acetate solutions (10.0 mM) were prepared in MilliQ water. Tris buffered saline (Tris 50.0 mM, NaCl 50.0 mM, pH 7.4) was prepared for spectroscopic experiments.

The copper-cuprizone complex has been previously characterized and is known to have an absorbance peak at 595 nm. An Agilent 8453 spectrophotometer was used to monitor the binding of copper to cuprizone. All samples were blanked with Tris buffer. For saturation curve experiments, final molarity of cuprizone was 8.8 x 10^-5^ M, and copper concentration was 2.2 x 10^-5^ M, 4.4 x 10^-5^ M, and 6.6 x 10^-5^ M to show the saturation of cuprizone in the cell. Saturation curves were produced in the presence of CNM-Au8 (26 μg/mL). After adding copper reagents, the samples were incubated for 5 minutes to stabilize the Cu-CPZ complex, and the full spectrum for each sample was recorded.

### Rat spinal cord culture and immunohistochemistry

A standard laboratory protocol was followed; in brief, postnatal rat spinal cord cells were dissociated and plated on PLL coated coverslips. Cells were allowed to adhere for at least 24 hours in base medium and then switched to vehicle or CNM-Au8 containing medium (final concentration of 0.01 ug/mL or 10 ug/mL), grown for 72 hours, fixed, and labeled with antibodies as follows: anti-Olig2, anti-MBP, anti-GFAP, anti-CC1.

## Acknowledgments

The consistent and enthusiastic support from Clene Nanomedicine’s CEO Rob Etherington, the support from our former CMO’s Drs. William Houghton, Consuelo Glenn, and Glen Frick, and the dedication of our amazing employees at Clene are greatly valued and appreciated.

We thank Pieter D’Arnaud of Instat, Inc. for help with statistical analyses of the cuprizone remyelination and fine motor kinetic data. We thank Dr. Damiano Fantini, Matthew J. Schipma and the staff at the NuSeq core (Northwestern University) for sequencing the RNAseq libraries and preliminary statistical analysis of sequence data (test statistic, p-value, false discovery rate), and especially Dr. Fantini for preliminary bioinformatics analyses of this data. We also thank Carol Cooke of the Electron Microscopy Lab in the Neurology Department of Johns Hopkins University for technical help with transmission electron microscopy. Cellomet, Neuro/sys, and Charles River Labs performed contracted research for us. Funding: This work was funded by Clene Nanomedicine, Inc.

## Author Contributions

AR and HET conducted the early and late cuprizone experiments and flow cytometry experiments. AR performed Seahorse experiments. AR and HET conducted the transcriptomics study. JZZ conducted the cuprizone pilot and replication studies. MM, ARD, and MGM produced CNM-Au8 and conducted the analytical and nanocatalytic characterizations of CNM-Au8. MK and RM performed all lysolecithin experiments as well as the isolated rat spinal OPC marker expression study. SK interpreted and qualitatively evaluated the corpus callosum micrographs from the cuprizone studies. MTH, SM, RM, MGM, and KSH designed the studies and analyzed data. KSH wrote the manuscript.

